# Alternative splicing and gene expression play contrasting roles in the parallel phenotypic evolution of a salmonid fish

**DOI:** 10.1101/2020.05.11.087973

**Authors:** Arne Jacobs, Kathryn R. Elmer

## Abstract

Understanding the contribution of different molecular processes to the evolution and development of divergent phenotypes is crucial for identifying the molecular routes of rapid adaptation. Here, we used RNA-seq data to compare patterns of alternative splicing and differential gene expression in a case of parallel adaptive evolution, the replicated postglacial divergence of the salmonid fish Arctic charr (*Salvelinus alpinus*) into benthic and pelagic ecotypes across multiple independent lakes.

We found that genes that were differentially spliced and differentially expressed between the benthic and pelagic ecotypes were mostly independent (<6% overlap) and were involved in different processes. Differentially spliced genes were primarily enriched for muscle development and functioning, while differentially expressed genes were mostly involved in energy metabolism, immunity and growth. Together, these likely explain different axes of divergence between ecotypes in swimming performance and activity. Furthermore, we found that alternative splicing and gene expression are mostly controlled by independent cis-regulatory quantitative trait loci (<3.4% overlap). Cis-regulatory regions were associated with the parallel divergence in splicing (16.5% of intron clusters) and expression (6.7 - 10.1% of differentially expressed genes), indicating shared regulatory variation across ecotype pairs. Contrary to theoretical expectation, we found that differentially spliced genes tended to be highly central in regulatory networks (‘hub genes’) and were annotated to significantly more gene ontology terms compared to non-differentially spliced genes, consistent with a higher level of connectivity and pleiotropy.

Together, our results suggest that the concerted regulation of alternative splicing and differential gene expression through different regulatory regions leads to the divergence of complementary phenotypes important for local adaptation. This study provides novel insights into the importance of contrasting but putatively complementary molecular processes for rapid and parallel adaptive evolution.

## Introduction

Since Turesson first used the term ‘ecotype’ nearly a century ago to describe genetically distinct populations that are adapted to alternative ecological environments (Turesson 1922), we have gained substantial insights into the genetics of ecological adaptation. Yet our knowledge of the molecular and regulatory mechanisms linking environmental influences, functional variation, and the development and evolution of new phenotypes in nature is still limited (Lewis and Reed 2019; Verta and Jones 2019). Divergence in gene expression has been strongly implicated in the rapid evolution and development of adaptive and divergent phenotypes, particularly as *cis*-regulatory mutations are thought to exhibit fewer deleterious effects than protein-coding changes (Prud’homme et al. 2007; Filteau et al. 2013; Manousaki et al. 2013; Alvarez et al. 2015; Campbell-Staton et al. 2017; Mack et al. 2018; McGirr and Martin 2018; Jacobs et al. 2019; Verta and Jones 2019). However, post-transcriptional processes, which have been suggested to play a substantial role in generating phenotypic variation (Li et al. 2016; Bush et al. 2017; Howes et al. 2017; Singh et al. 2017; Parenteau et al. 2019), have rarely been evaluated in cases of rapid adaptation in natural systems (Howes et al. 2017; Mallarino et al. 2017; Singh et al. 2017; Wang et al. 2018). This raises an important knowledge gap regarding the contribution and interaction of different molecular mechanisms in the evolution of ecologically adaptive phenotypes.

The alternative splicing of pre-mRNA transcripts is a post-transcriptional process that has been associated with phenotypic diversification in eukaryotes (Bush et al. 2017). Alternative splicing leads to the formation of distinct transcripts (‘isoforms’) through the retention or removal of different exons and introns from the immature mRNA of a single gene (Stamm et al. 2005; Nilsen and Graveley 2010; Bush et al. 2017). These isoforms can either encode structurally distinct proteins and result in the functional diversification of the proteome (Nilsen and Graveley 2010; Bush et al. 2017), or alter the regulation of transcript abundance through the formation of nonsense transcripts, which will be efficiently removed (Aznarez et al. 2018; Grantham and Brisson 2018). In eukaryotes, the majority of genes undergo splicing at some point during development (Grau-Bové et al. 2018). Splicing occurs through a dynamic ribonucleoprotein complex, the spliceosome (Lee and Rio 2015) and studies in model organisms showed that the expression and splicing of genes are mostly controlled by different genetic loci, indicating these processes can evolve independently (Li et al. 2016). Patterns of alternative splicing have been found to differ between closely-related species (Harr and Turner 2010; Singh et al. 2017) and the differential splicing of individual genes has in some cases been linked to the rapid evolution of ecologically-relevant phenotypic traits (Howes et al. 2017; Mallarino et al. 2017). In principle, alternative splicing and gene expression can evolve independently, either playing contrasting roles by affecting different genes and biological pathways (Grantham and Brisson 2018), or playing redundant roles by affecting the same genes (Singh et al. 2017; Healy and Schulte 2019). Despite our growing knowledge of the importance of alternative splicing in phenotypic evolution, the role of differential splicing has rarely been studied for cases of rapid ecological adaptation in natural systems, restricting our understanding of its influence and consistency in adaptive phenotypic diversification.

To address this gap, here we assess the pattern of alternative splicing associated with rapid phenotypic divergence in a natural and replicated system of ecological speciation. Arctic charr (*Salvelinus alpinus*) is a salmonid fish with a holarctic distribution and numerous and well-studied cases of independent parallel divergence into ecotypes, which differ in a range of ecological and phenotypic aspects along the depth axis within freshwater lakes (Jonsson and Jonsson 2001; Elmer and Meyer 2011; Elmer 2016). Benthic ecotypes mostly occupy the profundal or littoral habitat, show lower swimming activity, and usually have larger eyes and deeper and more robust heads, allowing them to better handle benthic prey (Jonsson and Jonsson 2001,Adams and Huntingford 2002b; Alekseyev et al. 2002; Klemetsen 2002). In contrast, pelagic ecotypes occupy the open water where they forage for plankton and are thus more active swimmers, with mostly smaller and more slender heads (Jonsson and Jonsson 2001,Adams and Huntingford 2002b; Alekseyev et al. 2002; Klemetsen 2002). These divergent ecotypes have evolved independently following the last glacial maximum 10,000 – 15,000 years ago (Jacobs et al. 2020). Patterns of genomic differentiation and evolutionary histories differ widely across Arctic charr ecotype pairs, while patterns of gene expression divergence are more predictable and consistent across populations, evolution across distinct phylogeographic lineages (Guðbrandsson et al. 2018; Jacobs et al. 2020). Both heritable genetic and plastic changes have been shown to contribute to this ecotype divergence (Adams and Huntingford 2002a, 2004; Klemetsen 2002; Garduno-Paz et al. 2012; Jacobs et al. 2020).

We investigated the patterns and contributions of differential gene expression, alternative splicing and genomic changes to rapid and parallel eco-morphological divergence in these natural populations. To do so, we reanalysed RNA-seq data (Jacobs et al. 2020) generated from white-muscle from three Arctic charr ecotype pairs across three independent lakes (Fig. 1a, Table 1). We investigated genome-wide gene expression, signals of selection and used two complementary approaches to study patterns of alternative splicing. This provides novel insights into the alternative roles of different molecular processes underlying rapid ecological and phenotypic divergence in environmental context and fills an important knowledge gap in our understanding of the link between genotype, environment and adaptive phenotypes in natural populations.

**Fig. 1.**
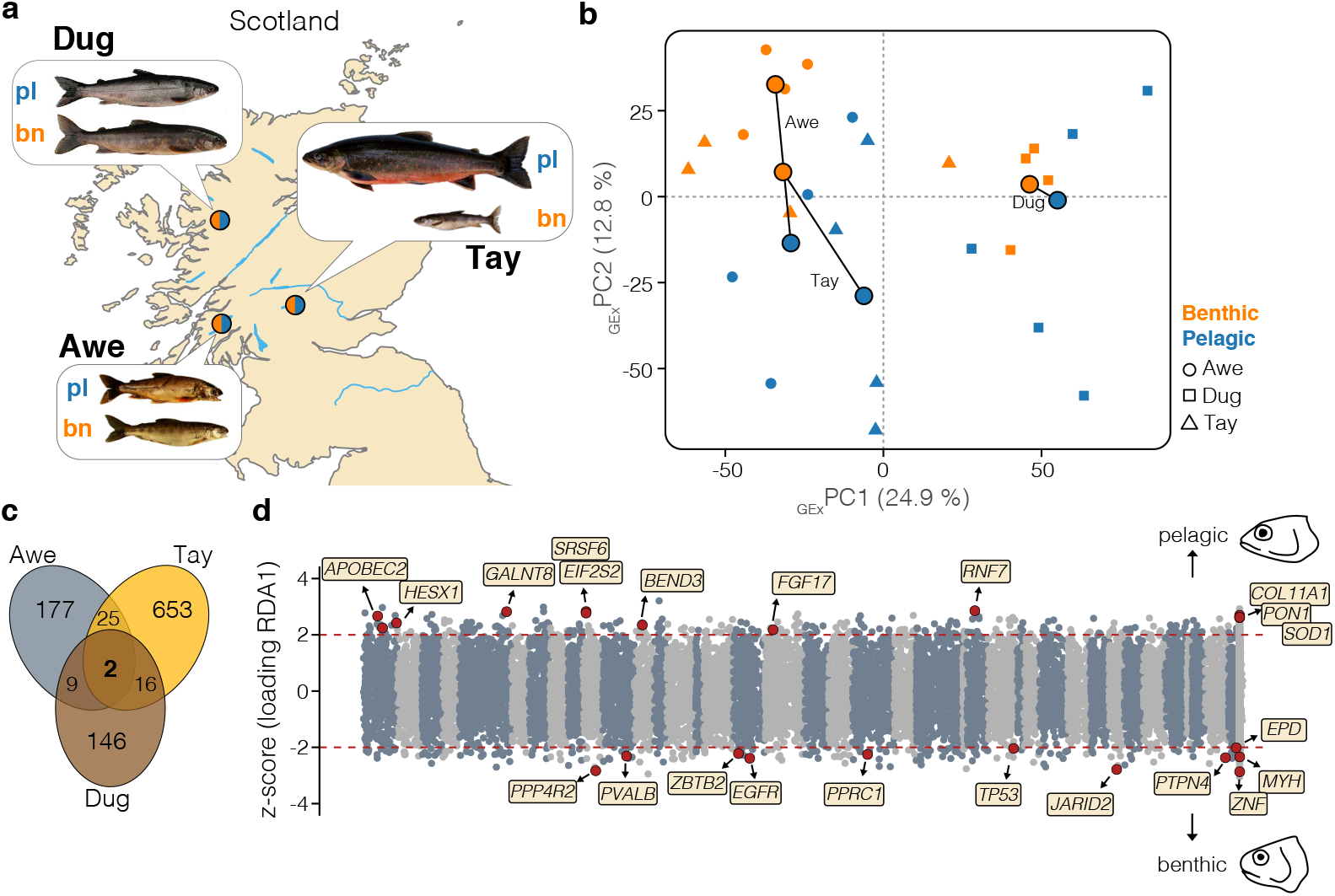
Sampling and differential gene expression. a) Map showing the locations of the focal populations (Awe, Tay and Dughaill [Dug]). Representative pictures of the benthic and pelagic ecotypes per lake are shown. Coordinates and additional sample information are given in Table 1. b) PCA based on expression data (GEx-PCA) of all genes (n=19,623) for all individuals (n=24). Round large points show the respective centroids for each ecotype, and sympatric ecotypes are connected by vectors. Small unicolour points represent individuals. Populations are coded by shape and ecotypes by colour. c) Venn diagram displaying the amount of overlap of differentially expressed genes between sympatric ecotypes (FDR < 0.05) across lakes. d) Distribution of z-scores derived from the RDA, depicting association of gene expression with ecotype across the genome. The red dashed lines highlight the significance threshold (|z| > 2). Genes with negative z-scores were associated with the benthic ecotype across lakes, and genes with positive thresholds were associated with pelagic ecotypes. Illustrations highlight differences in head shape between benthic and pelagic ecotypes. Core genes that were differentially expressed in at least two ecotype pairs and significantly associated with ecotype across lakes are highlighted in red, with gene IDs from the Arctic charr genome annotation. Chromosomes are highlighted by alternative colours, and unplaced scaffolds are located at the end.

**Table 1.** Study population summaries.

| Population | Lat.(N)/Lon.(W) | Ecotypes | Evol. History<sup>1</sup> | DE | DS (DIE/DEU) | Div. SNPs |
| --- | --- | --- | --- | --- | --- | --- |
| Awe | 56°20'/005°05' | Bn - Pl | IM | 209 | 1,475 (85/1428) | 471 |
| Tay | 56°30'/004°10' | Bn - Pl | SC | 692 | 197 (48/164) | 357 |
| Dughaill | 57°28'/005°20' | Bn - Pl | SC | 167 | 148 (106/67) | 396 |

*Note: Ecotypes: Benthic (Bn) and Pelagic (Pl); Evolutionary histories: likely evolved under isolation-with-migration (IM), likely evolved under secondary contact (SC); DE - Differentially expressed gene; DS - Differentially spliced gene (based on differential intron excision (DIE) and differential exon usage (DEU); Div. SNPs - Number of divergent SNPs showing signs of selection based on Rsb score. [1] Jacobs et al. 2020.*

## Material and Methods

### Data

RNA-seq data were drawn from (Jacobs et al. 2020) (NCBI BioProject: PRJNA551374). Briefly, adult Arctic charr had been sampled from Loch Awe, Loch Tay and Loch Dughaill in Scotland (Fig. 1a, Table 1) during spawning time. High quality RNA was extracted from white muscle tissue and RNA-seq libraries were prepared for 24 individuals (n=4 per ecotype per lake) using the TruSeq Stranded mRNA Sample Preparation kit. Libraries were paired-end sequenced to an average depth of 25-30M reads per library.

### Filtering and read mapping

Adapters and low-quality reads were trimmed using *Trimmomatic* v0.36 and reads were aligned against the Arctic charr reference genome (ASM291031v2) (Christensen et al. 2018) with *STAR v2.5.2b* using a two-step mapping approach and duplicates were marked using the *picard* tool.

### Gene expression analyses

Raw reads for each transcript were counted using *HTSEQ-count*, and subsequently filtered, normalized and analyzed using *DESeq2*. Transcripts with <20 reads across all samples were excluded. Principal component analysis (PCA) was performed based on rlog-transformed read counts using *pcaMethods* (R-package) with the following settings: scaling = “none”, center = TRUE. First, we identified differentially expressed genes between benthic and pelagic ecotypes using a pairwise analysis by lake in *DESeq2*. Second, we used a conditioned redundancy analysis (*RDA*) in *vegan* (R-package) to identify genes with expression patterns associated with ecotype (binary), while correcting for lake effect. We selected genes with z-transformed loadings above 2 or below −2 as associated with ecotype. Furthermore, we constructed gene co-expression networks using *WGCNA* (R-package) based on *rlog*-transformed read counts.

Modules were defined using the dynamic treecut algorithm, with a minimum module size of 25 genes and a cut height of 0.992, and similar modules were merged using an eigengene distance threshold of 0.25. We assessed module-trait correlations for lake, ecotype and sex. We further estimated the scaled intramodular connectivity (k_within_) for all genes within modules, excluding unassigned genes. We compared k_within_ between candidate genes (differentially spliced or expressed) and all other expressed genes using Wilcoxon rank sum tests.

### Alternative splicing analysis

First, we used *DEXseq* to analyse differential exon usage between ecotypes while taking biological replicates into account. We used the modified *HTSeq-count* script provided with *DEXseq* to quantify exon-specific read counts for each sample. Differential exon usage was estimated per lake using the following model structure: ~ sample + exon + ecotype:exon. A PCA based on rlog-transformed exon expression counts was performed in *pcaMethods*. Second, we used *LeafCutter* (Li et al. 2018) to perform intron clustering and differential splicing analyses for each lake separately (minimum coverage of 10 reads per intron, minimum of 4 samples supporting an intron, minimum of 2 samples per ecotype supporting an intron). This method is independent of the genome annotation. Differential splicing of intron clusters was measured as the ‘change of percent spliced in’ (ΔPSI). We used a combined dataset for visualising splicing patterns within and across lakes using a PCA based on intron cluster count ratios and sashimi-plots using the *LeafViz* shiny-app that is distributed with *LeafCutter*.

### SNP calling, SNP effect prediction and signatures of selection

We called single nucleotide polymorphisms (SNPs) from the genome aligned RNA-seq data using *freebayes*, after marking duplicates using *picard*, using coverage threshold of three. We filtered the biallelic SNP dataset using the *vcffilter* command in *vcflib* and *vcftools*, retaining biallelic SNPs with i) phred quality score above 30, ii) genotype quality above 20, iii) an allele-depth balance between 0.25 and 0.75. We furthermore filtered for Hardy-Weinberg disequilibrium (p-value threshold < 0.01) and only kept sites that were present in at least 90% of all individuals across populations. We annotated the retained SNPs and predicted their effect, particularly splice-site disrupting variants, with *SnpEff*. PCA was based on LD-pruned SNPs in *SNPrelate* (R-package). We identified SNPs putatively under selection by comparing patterns of extended haplotype homozygosity (Rsb score) between sympatric ecotypes using *rehh* (R-package). We identified haplotypes under selection as those with absolute Rsb values above 4. Phasing and imputation were performed with *fastPhase*.

### Expression and splicing QTL mapping

We identified genetic variants putatively underlying variation in gene expression, we used an expression quantitative trait locus (*cis*-eQTL) approach to identify *cis*-acting variants associated with expression. We used linear models with lake as a covariate implemented in *MatrixEQTL* v.2.2 to associate SNPs with the expression of nearby genes (< 1Mb) and a false-discovery rate below 0.05. Using the same approach, we also tested for cis-regulatory splicing QTL (*cis*-sQTL) based on association with intron excision ratios from *LeafCutter*.

### Functional gene ontology analysis

We used the gene ontology (GO) annotation provided with the published Arctic charr reference genome (Christensen et al. 2018) as the basis for analyses of biological processes and molecular functions. We used *Blast2Go v.5.2.4* to perform overrepresentation analysis using Fisher’s Exact tests and gene set enrichment analysis (GSEA). We clustered GO terms using the *revigo* clustering algorithm to account for redundancy, and visualised similarities between clustered GO terms using multidimensional scaling (MDS) scaling plots based on the semantic similarities of GO, as well as interaction networks in *Cytoscape*.

### Statistical analysis of sharing across lakes

We performed hypergeometric tests using the *phyper* R-function to calculate the probability that differentially regulated genes are shared more or less often across two lakes than expected by chance. To identify if more or fewer genes were shared than expected, we calculated a representation factor (RF), which compared the observed number shared genes to the expected number (e.g. [Number of DS x Number of DE]/all expressed genes). An RF > 1 indicates that more genes than expected are shared, while an RF < 1 indicates that fewer than expected are shared (Grantham and Brisson 2018).

## Results

### Divergence in gene expression between ecotypes

We investigated patterns of gene expression divergence between sympatric Arctic charr ecotypes within and across the three lakes (Fig. 1a, Table 1). PCA based on gene expression data for 19,623 transcripts showed that the lake of origin had the strongest effect on expression patterns (PC1=24.9%; linear model [LM] effect size for PC1: *η*^2^_Lake_=0.816, P<0.001; Fig. 1b), consistent with the independent divergence history of ecotype pairs since the last glacial maximum (Jacobs et al. 2020). Benthic and pelagic ecotypes separated along PC2 across lakes (12.8%; LM: *η*^2^_Ecotype_=0.296, P=0.013) (Fig. 1b), indicating consistent patterns of gene expression divergence. Parallelism was further supported by nonsignificant ‘ecotype x lake interaction’ terms for PC1 (LM: *η*^2^_Ecotype x Lake_=0.064, P=0.552) and PC2 (LM: *η*^2^_Ecotype x Lake_=0.136, P=0.269), the interaction term representing non-parallel differences in gene expression across lakes.

We detected between 169 and 692 differentially expressed (DE) genes between sympatric ecotypes (Fig. 1c, Table 1, Fig. S1, Table S1), with ecotype pairs from two different lakes sharing more DE genes than expected by chance (Hypergeometric tests [HGT]; P < 0.001, RF between 2.7 and 5.0; RF > 1 indicates more overlap than expected). Combined analyses of expression across all samples and lakes identified 699 genes that showed ecotype-associated expression, with ecotype explaining 6.8% of gene expression variation (ANOVA: F_(1,20)_=1.914, P=0.007) (Fig. 1d, Figure S1b). Of these 699 ecotype-associated genes, 23 were differentially expressed in at least two of the three populations (Table S1).

### Patterns of differential splicing within and across lakes

We investigated alternative splicing patterns within and across ecotype pairs using two independent approaches (see methods). The lake of origin explained the largest proportion of variance in splicing based on intron excision ratios (n=18,207 intron-clusters; DIE) (PC1: LM: *η*^2^_lake_=0.878, P=5.97e-09; Fig. 2a). The major axes of variation in splicing also strongly separated ecotypes (Fig. 2a), with ecotype explaining a substantial proportion of variance along PC1 and PC2 (PC1 [8.4%]: *η*^2^_Ecotype_=0.729, P=1.67e-06; PC2 [7.1%]: *η*^2^_Ecotype_=0.593, P=7.21e-05;). The non-parallel ‘lake x ecotype’ interaction effect explained a small but significant proportion along PC1 (LM: *η*^2^_Ecotype x Lake_=0.456, P=0.0042), but not along PC2 (*η*^2^_Ecotype x Lake_=0.272, P=0.058), indicating variation in alternative splicing patterns across ecotype pairs. The PCA based on exon usage (DEU; Fig. S2a) was similar to the gene expression PCA.

**Fig. 2.**
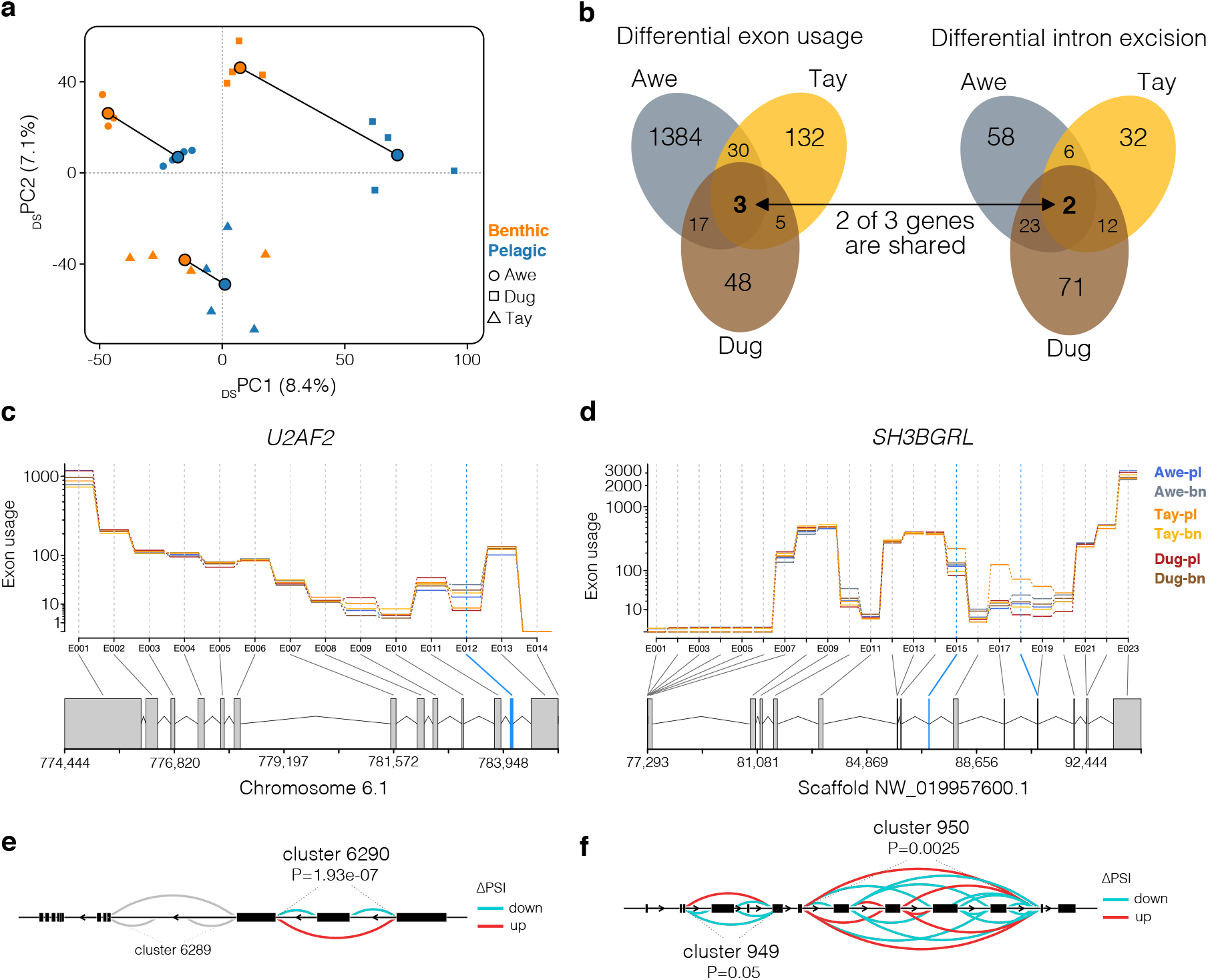
Shared patterns of differential splicing across lakes. a) PCA based on intron excision ratios (n=18,207 introns) for all individuals. See Fig. 1 for explanation of the legend. b) Venn diagrams showing the amount of overlap of differentially spliced genes based on differential exon usage and differential intron excision between sympatric ecotypes across lakes. Two genes were detected using both approaches. c,d) Gene models illustrating alternative splicing patterns for c) U2AF2 and d) SH3BGRL based on exon usage. The expression of each exon corrected for overall gene expression (exon usage) is shown for each ecotype. Exons that are differentially spliced in at least two of the three ecotypes are highlighted in blue. e,f) Sashimi graphs highlighting patterns of differential intron excision (DIE) across all ecotypes and lakes for intron clusters associated with e) U2AF2 and f) SH3BGRL. The amount of DIE is measured as ‘change in the percent spliced in (ΔPSI)’.

Analysing differential splicing between sympatric ecotypes, we found between 48 and 104 genes to be associated with differentially spliced intron clusters, and between 67 and 1428 genes to show differential exon usage (in one or several exons) (Fig. 2b, Table 1), corresponding to about 1 to 8% of all expressed genes. Between 1.2 and 27.1% of all differentially spliced genes (5 to 30 DS genes) were detected in at least two of the three ecotype pairs (Fig. 3a, Fig. S2 and S3, Table S2; HGTs; all P<0.05, RF between 33.9 and 51 for DIE; RF between 2.7 and 12.6 for DEU). Two genes were differentially spliced in all three lakes, based on DEU and DIE (*U2AF2, SH3BGRL*; Fig. 3b-e; Fig. S3a,b). Additionally, *FTH1* (Fig. S3c) was differentially spliced based on DEU in all three lakes. The overall proportion of expressed genes that were differentially spliced was on average 2.82 ± 3.87% (mean ± SE) based on DEU and 0.40 ± 0.15% based on DIE.

**Fig. 3.**
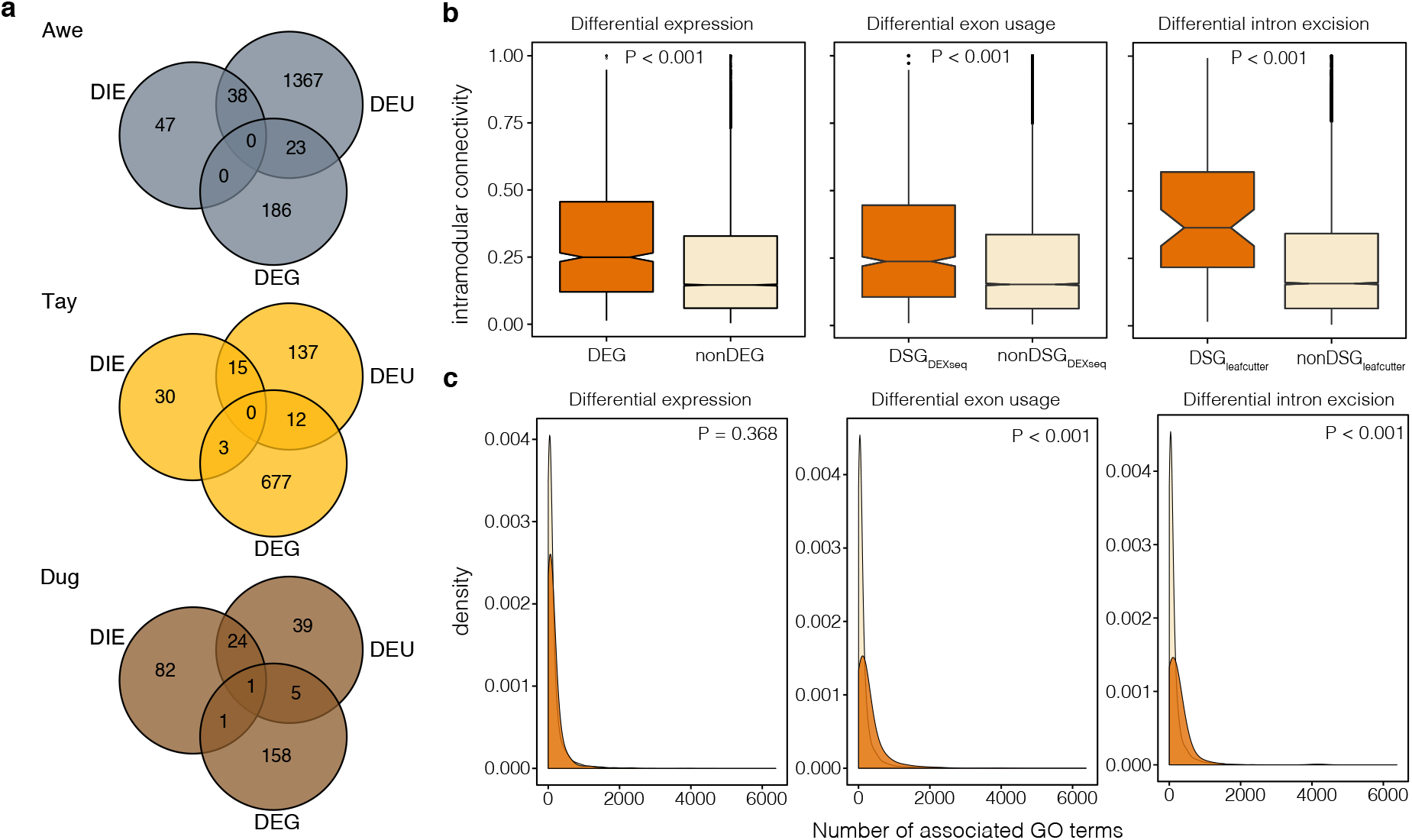
Sharing, connectivity and pleiotropy of differentially spliced and differentially expressed genes. a) Venn diagrams showing the amount of overlap of differentially spliced (by analyses) and differentially expressed genes for each lake. b) Boxplots (bar = median; notch = confidence interval around the median, box range = range between third and first quartile (interquartile range); whiskers = extend to furthest point (highest or lowest) no further than 1.5 times the interquartile range; points = outliers) showing differences in intramodular connectivity between candidate and non-candidate genes for DEGs (n=964 vs n=15391), DEU (n=926 vs n=16355) and DIE (n=103 vs n=16355). c) Differences in the number of associated gene ontology (GO) terms (biological processes) for candidate genes and noncandidate genes, based on DEGs and DSGs (DEU and DIE). DS genes were associated with more GO terms compared to the genomic background (non-candidate genes), suggesting higher pleiotropic effects.

Comparing splicing events across ecotype pairs, we found that in some cases different intron clusters were differentially spliced within the same genes, e.g. in the two highly shared DS genes *U2AF2* and *SH3BGRL* (Fig. S3a,b). Similar results were detected based on different exon usage (Fig. S3c-e). For example, while benthic ecotypes consistently displayed a significantly higher usage of exon 12 compared to pelagic ecotypes in *U2AF2* (Fig. S3c), exons 1, 7 and 11 were only differentially spliced between ecotypes in Awe (Fig. S3c). This suggests that differential splicing is parallel at the gene level, but that isoform diversity likely differs across lakes.

### Alternative splicing and differential gene expression are functionally non-redundant

We found that DS genes were generally not differentially expressed, with 5.6 ± 4.1% of DS genes being differentially expressed within ecotype pairs (1.7% in Awe, 5.2% in Dug to 9.9% in Tay; Fig. 3a). While DS genes identified based on DIE were not differentially expressed (HGT: P > 0.05, RF between 0 and 1.77), DS genes based on DEU tended to be differentially expressed more often than expected by chance in all three ecotype pairs (HGT: P < 0.05, RF between 1.5 and 8.7). However, this overrepresentation was very low and could stem from technical artefacts, namely that both *DEXseq* (DEU) and *DEseq2* (DE) analyses rely on read count data of annotated features (Anders et al. 2012).

### Increased connectivity and pleiotropy of candidate genes

To better understand the regulatory context and putative importance of DS and DE genes, we investigated gene coexpression networks (Langfelder and Horvath 2008) and pleiotropy based on gene ontology annotations (McGirr and Martin 2018). We found that both DE and DS genes showed higher degrees of connectivity than non-DE (Wilcoxon rank sum test: P < 0.001) or non-DS genes (Wilcoxon rank sum test: P < 0.001) (Fig. 3b, Fig. S5) in gene co-expression networks, consistent with more central positions of differentially regulated genes in regulatory expression networks.

DS genes were on average associated with more gene ontology terms (biological processes) than non-DS genes (Fig. 3c), indicating a greater level of pleiotropy. DS genes were on average annotated with 265 (s.d = 412) and 194 (s.d = 369) GO terms for DEU (*DEXseq*) and DIE (*leafcutter*), respectively, whereas non-DS genes were annotated with an average of 142 (s.d = 264) and 141 (s.d = 263) GO terms (Wilcoxon rank sum tests; DEU: P < 0.001; DIE: P < 0.001). We did not find any difference in the number of annotated GO terms between DE (mean = 141, s.d = 245) and non-DE genes (mean = 148, s.d = 289; Wilcoxon rank sum test: P = 0.368; Fig. 3c). Together, our results suggest that DS genes, and to a lesser degree DE genes, hold central functional roles in regulatory networks and are more pleiotropic.

### Different regulatory processes affect different biological pathways

We compared functional annotations (gene ontology terms) of genes with ecotype-associated gene expression (RDA analyses) and DS genes, to identify putative functional downstream differences. GO terms that were significantly enriched for ecotype-associated DE genes (RDA analyses) in a GSEA were related to metabolic processes (e.g. oxidative phosphorylation) and translational activity (e.g. cytoplasmic translation) in pelagic ecotypes, and cell growth and differentiation (e.g. lymphocyte differentiation), immune response (e.g. interleukin-1 secretion) and signal transduction (e.g. positive regulation of endocytosis) in benthic ecotypes (Fig. 5a, Table S3). Similar processes and functions were identified using a GO term overrepresentation analysis of all significant ecotype-associated DE genes, although none of the GO terms was significantly overrepresented after correcting for multiple testing (Table S4). In contrast, overrepresentation analysis of DS genes revealed an enrichment of biological processes and molecular functions related to muscle development and functioning (Fig. 5b, Table S5), such as myofibril assembly, actinin binding, muscle structure development. Other processes enriched for DS genes included RNA-binding processes, mRNA splicing and translational regulation (Fig. 5b, Table S3-5). GO terms that were enriched for DE and DS genes were mostly involved in gene regulation, such as translation, transcription and RNA catabolism (Fig. 5c, Fig. S6). The proportions of overlap were relatively low, with an average of 13.0% (range from 0 to 25.8%) of DS-enriched GO terms overlapping DE-enriched GO terms (Fig. S7). Overall, we find that DS and DE genes in white muscle tend to be involved in different molecular functions and biological processes.

### Genetic variation underlying gene regulation

Lastly, we mapped the genetic basis of gene regulation and identified genetic signatures of selection to assess their impact on regulatory evolution and redundancy. We identified 101,487 high-confidence SNPs (Fig. S6), of which 2,404 (located in 2106 genes) were predicted to have effects on splicing using *snpeff*. Genes containing high-impact splice-site variants were more likely to show differential intron excision (5.2% of all DS-DIE genes; HGT: P = 8.54e-15). We further identified 1,919 *cis*-sQTL (*cis*-regulatory splicing quantitative trait loci) associated with variation in intron excision ratios of 626 intron clusters across all the three lake populations (FDR < 0.05, Fig. 5a). Of these, five *cis*-sQTL were predicted to be high-impact splice variants, and 40 intron clusters associated with *cis*-sQTL were also differentially spliced (HGT: P = 0.00013, RF = 22; expected intron clusters = 21; Table S6).

Furthermore, we identified 1,562 *cis*-eQTL (cis-regulatory expression quantitative trait loci) associated with the expression variation of 734 genes (FDR < 0.05, Fig. 5a), with between 6.7 and 10.1% of DE genes being associated with *cis*-eQTL (Fig. 5c, Table S7). We found 117 *cis*-sQTL and *cis*-eQTL to be shared (HGT: P = 6.14e^-08^, RF = 70.4; expected number of shared QTL = 67), suggesting some genomic regions regulate both splicing and expression in Arctic charr. However, the majority of cis-regulatory regions (96.64%) only affected one of the two regulatory processes. Overall, this suggests that differential splicing and expression are both at least in part genetically regulated, most likely by different genomic regions.

We further tested if differentially regulated genes were associated with genetic signatures of selection (Table S8). Neither DE nor DS genes were significantly associated with genetic signatures of selection (HGT: all P > 0.5). Only two *cis*-sQTL and two *cis*-eQTL showed signs of selection in at least one lake. However, SNPs putatively under selection in at least one population were more often located in the 3’ untranslated region (3’-UTR) compared to the full SNP dataset (20.7% of selected SNPs vs. 8.98% of all SNPs; *χ*^2^ = 4.53, P = 0.033) and were significantly more often synonymous mutations (17.94% of selected SNPs vs. 7.17% of all SNPs; *χ*^2^ = 4.35, P = 0.037) (Fig. 5d). Thus, putatively selected SNPs might underlie gene expression changes between ecotypes through changes of *cis*-regulatory elements in the 3’-UTR rather than structural protein changes.

## Discussion

Post-transcriptional processes are widely understudied in evolutionary and ecological genomics of natural populations, leading to a knowledge gap in our understanding of their importance for adaptation. Here, we analysed white-muscle RNA-seq data from three replicated benthic-pelagic Arctic charr ecotype pairs to contrast the patterns and regulation of alternative splicing and gene expression. We found evidence suggesting that alternative splicing and differential expression, which are both highly parallel across lakes, play contrasting but complementary functional roles by affecting different genes and pathways. These processes are likely regulated through alternative genetic changes and plastic responses, as evidenced by the integration of expression and splicing QTL mapping with selection analyses. Differential splicing between ecotypes potentially facilitated the rapid adaptive evolution of Arctic charr, due to its central position in regulatory networks and increased pleiotropy. Overall, this study provides novel insights into the molecular processes underlying adaptive parallel divergence and suggests alternative roles for complementary regulatory processes in rapid and parallel adaptive diversification.

### Alternative splicing and gene expression have contrasting phenotypic effects

Contrary to other studies of rapid adaptation or acclimation in teleosts (Singh et al. 2017; Healy and Schulte 2019), we found that most DS genes were not differentially expressed between ecotypes (Fig. 3a). This lack of overlap suggests that these processes might be independently regulated (Li et al. 2016) and potentially affect different biological pathways and phenotypes. Indeed, DS and DE genes were enriched for different biological processes and molecular functions (Fig. 4, Fig. S5). We found that DS genes were mostly involved in processes related to muscle development and function through the alternative splicing of genes with molecular functions involved in actinin binding (Fig. 4b). Functional differences in muscle formation and functioning are consistent with differences in swimming performance and activity between benthic and pelagic ecotypes that require adaptive changes in muscle composition and arrangement (Altringham and Ellerby 1999; Klemetsen 2002). In contrast, DE genes were mostly enriched for metabolic and developmental processes (e.g. mitochondrial respiratory chain complex, post-embryonic organ development) (Fig. 4a), consistent with putative differences in activity and metabolic requirements between ecotypes with different foraging and swimming habits (Klemetsen 2002; Evans and Bernatchez 2012; Dalziel et al. 2018). Metabolic processes in particular have been observed to be differentially regulated based on expression in other cases of eco-morphological diversification in teleosts (Filteau et al. 2013; McGirr and Martin 2018) and were found to be under selection in salmonid fishes (Schneider et al. 2019). Alternative splicing in vertebrates, however, has been suggested to diverge more rapidly than gene expression (Barbosa-Morais et al. 2012; Merkin et al. 2012), indicating that it might play a key role at early stages of divergence and ecological speciation. Thus, divergence in muscle development and function might have preceded metabolic divergences. Comparative studies across replicated ecotype pairs with different divergence times are needed to better understand the role of splicing in rapid eco-morphological divergence on postglacial timescales.

**Fig. 4.**
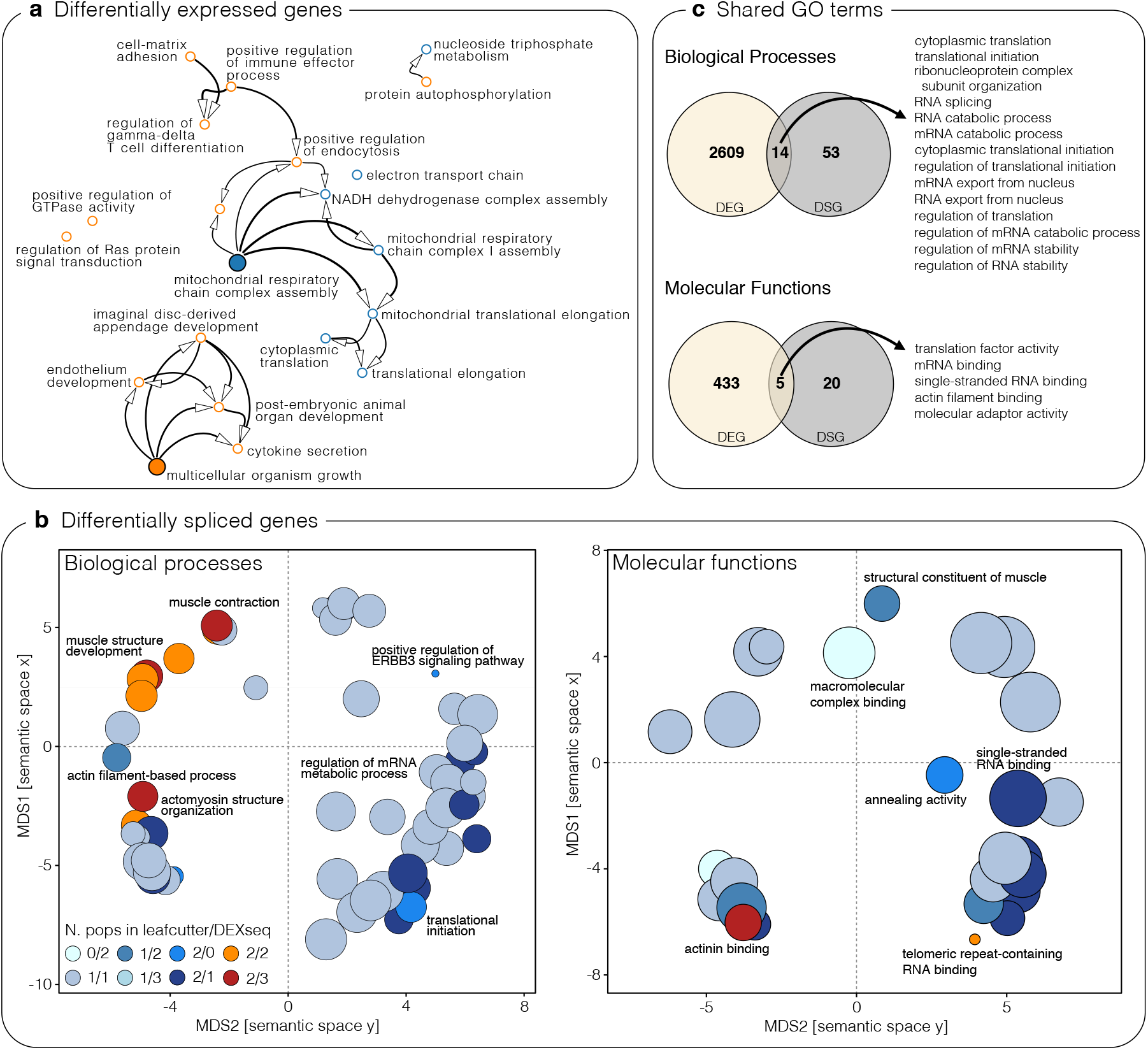
Divergent functional pathways. a) Gene ontology (GO) term interaction network for GO terms enriched for ecotype-associated genes (RDA). Benthic ecotype associations are highlighted in orange, while pelagic ecotype associations are blue. Central processes are highlighted by larger filled dots. b) Multidimensional scaling (MDS) plots for shared GO terms (Biological processes; Molecular functions) enriched for differentially spliced genes (DIE and DEU). Clustering was performed based on semantic similarity of GO terms. Circles are coloured based on the number of populations they were enriched in and if they were detected based on DIE (leafcutter), DEU (DEXseq) or both. The most representative and highly shared GO terms are named. The size of the circles corresponds to the number of genes associated with a GO term. c) Venn diagrams showing the overlap between GO terms (Biological processes, Molecular functions) associated with differentially spliced (DS) and differentially expressed (DE) genes. The names of overlapping processes and functions are listed.

Alternative splicing of transcripts can either lead to new functional proteins or to the regulation of transcript abundance through splicing-related nonsense-mediated decay (NMD) (Stamm et al. 2005). The role of NMD in regulating transcript abundance between ecotypes is supported by the differential splicing and regulation of *SRSF7* (Fig. 5b), a splicing factor that has been linked to regulating NMD pathway (Königs et al. 2020). In general, genes involved in the transcription and splicing machinery were both differentially expressed and spliced (Fig. 4c), suggesting that gene regulatory processes are highly divergent between ecotypes. This includes *U2AF2* on LG 6.1 (Fig. 2c,e), which is differentially spliced in all three ecotype pairs and encodes for a splicing factor important for pre-mRNA processing. Interestingly, we observed that a paralog of *U2AF2* on LG 33 was differentially expressed between ecotypes. This suggests that paralogs, which potentially originated through the whole-genome duplication in salmonids (Lien et al. 2016), might be regulated through different molecular processes, thus increasing functional diversity while minimising constraints (Bush et al. 2017; Iñiguez and Hernández 2017). A similar pattern was observed for *MYH*. However, more analyses are needed to test this pattern (Iñiguez and Hernández 2017) and assess the role of the whole-genome duplication in salmonids on the regulatory divergence between recently diverged ecotypes. Without detailed molecular studies of translation, protein abundance and transcript decay for candidate genes, we cannot discern the exact molecular and phenotypic impact of splicing and expression events (Mallarino et al. 2017). Overall, these findings are consistent with our hypothesis that alternative splicing and gene expression play contrasting functional roles in the divergence of Arctic charr ecotypes.

**Fig. 5.**
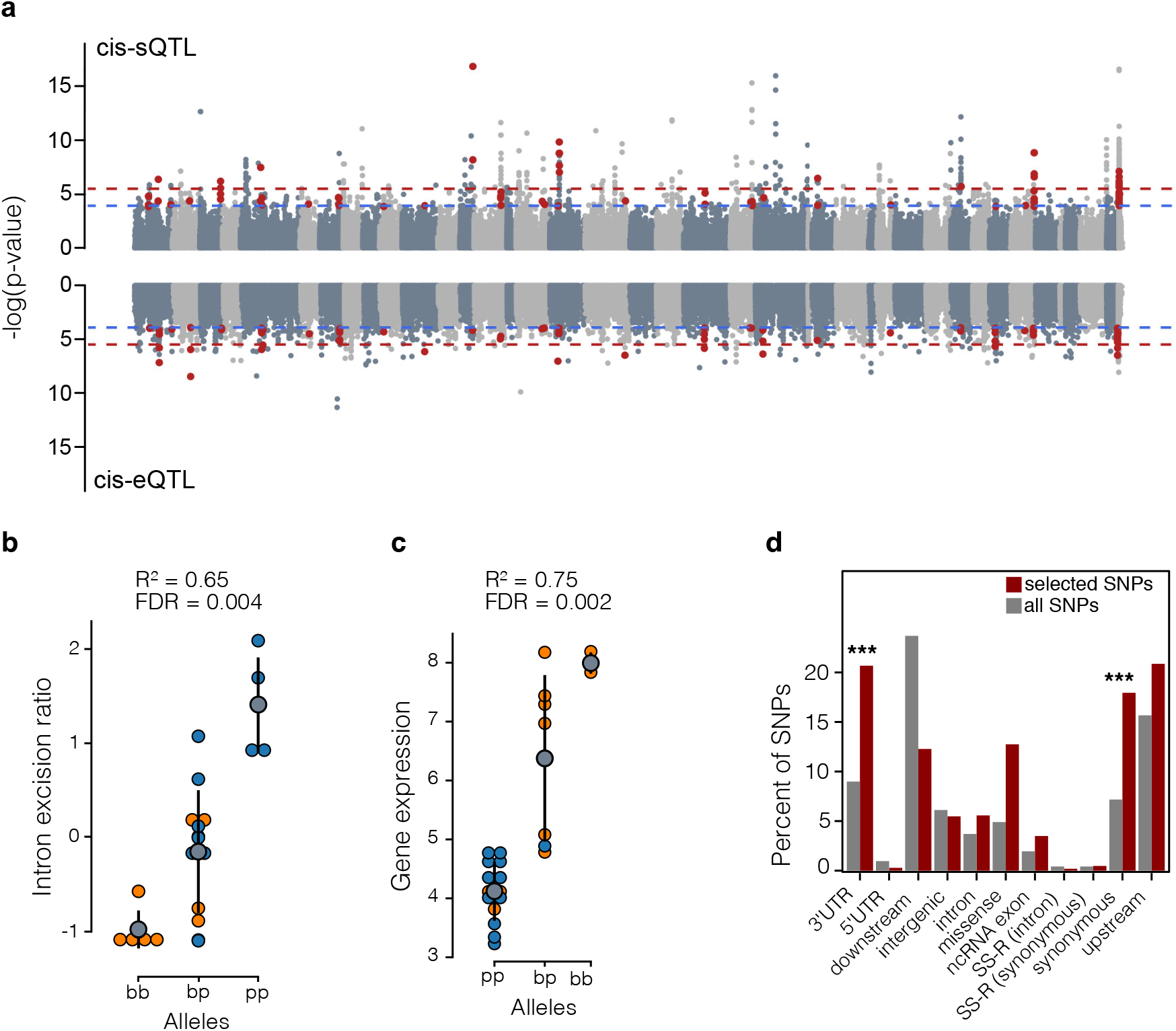
Genetic variation underlying regulatory variation. a) Manhattan plots showing the association of SNPs with variation in alternative splicing (intron excision ratios) across ecotypes and lakes (top) and with variation in gene expression (bottom). SNPs that are highlighted in red were detected in both analyses (n=117). Chromosomes are highlighted by alternative colours, and unplaced scaffolds are located at the end. The blue dashed line indicates a false-discovery rate (FDR) of 5% and the red dashed line an FDR of 1%. b) *cis-* sQTL associated with the intron excision of an intron cluster located in *SRSF7*. The y-axis shows the intron excision ratio for the intron cluster by genotype (p = pelagic allele, b = benthic allele) across individuals (points) and ecotypes (colour; orange = benthic, blue = pelagic). The grey dot and ranges show the mean intron excision ratio by genotype and the standard deviation. c) *cis*-eQTL associated with the normalized gene expression of *EGFR*. The plot shows how the expression of *EGFR* differs with genotype (p = pelagic allele, b = benthic allele) across individuals (points) and ecotypes (colour; orange = benthic, blue = pelagic). The grey dot and ranges show the mean expression per genotype and the standard deviation. Both *SRSF7* and *EGFR* are also differentially spliced or expressed in at least two ecotype pairs. d) Predicted effects and locations of SNPs under selection. A larger proportion of SNPs under selection (in at least one lake) are located in 3’-UTRs or are synonymous compared to proportions in the full SNP datasets (P < 0.001).

### Parallel divergence in alternative splicing across replicated ecotype pairs

While it is known that gene expression patterns are often highly parallel across ecotype pairs, even across lineages (Filteau et al. 2013; Manousaki et al. 2013; Rougeux et al. 2019; Jacobs et al. 2020), less is known about patterns of alternative splicing in ecological context. For example, parallel splicing has been described across cichlid species occupying parallel trophic niches (Singh et al. 2017), suggesting that alternative splicing might play a role in the evolution and development of replicated adaptive phenotypes. Similarly, we found highly parallel patterns of alternative splicing across the three replicated ecotype pairs (Fig. 2a), with dozens of transcripts being differentially spliced in two or all three ecotype pairs (Fig. 2b). This pervasive parallelism in splicing and expression on the gene level suggests strong regulatory constraints and low regulatory redundancy, e.g. due to limited standing regulatory variation across these populations (Yeaman 2015; Rougeux et al. 2019).

Despite parallelism in splicing on the gene level, we found that isoform usage differed in some cases across lakes (Fig. 2c-f). While the same genes were detected to be differentially spliced in all three ecotype pairs (*U2AF2, SH3BGRL* and *FTH1*), the exact differentially expressed exon (*DEXseq*) or excised intron cluster (*LeafCutter*) differed in some cases between lakes (Fig. S3). For example, in *U2AF2*, which encodes a splicing factor, the same intron cluster is differentially spliced in Awe and Tay (Fig. S3a). However, in Dughaill an overlapping but distinct intron cluster is differentially spliced (Fig. S3a), leading to the formation of a different isoform. Such subtle isoform differences can lead to differences in the function and/or fate of transcript, and can therefore have drastically different downstream effects (Keren et al. 2010; Eksi et al. 2013; Mallarino et al. 2017). Differences in isoform usage further suggest that the underlying genetic changes and/or splicing mechanisms differ across lakes, either due to different environmental impacts that alter splice-site choice or differences in the regulatory genetic architectures (Keren et al. 2010). The differential splicing of the same genes, despite the usage of different isoforms, indicates a potential functional role of these genes in the ecological divergence of Arctic charr, although the functional role of these isoforms might differ. Proteomic and functional analyses of these isoforms will be needed to better understand the downstream impacts of such splicing differences.

### Central regulatory roles and pleiotropy of alternatively spliced genes

The effect of differential regulation on phenotypic divergence has been suggested to be stronger for genes that are more central in regulatory networks (‘hub genes’) and show a higher degree of pleiotropy (Batada et al. 2006; Filteau et al. 2013), as simple changes to such central genes can have rapid and substantial phenotypic effects (Koubkova-Yu et al. 2018). On the other hand, it has been suggested that DS genes are less likely to be hub genes due to constraints of strong co-expression correlations and potentially increased pleiotropy (Iñiguez and Hernández 2017). However, we found that DS and DE genes were highly central in regulatory networks (Fig. 3b), which suggests that the differential regulation of these genes potentially has strong functional impacts on the divergence of Arctic charr. Constraints associated with the high complexity of altering pleiotropic or hub genes has led to the suggestion that they are less important for rapid adaptive evolution (Papakostas et al. 2014; Mäkinen et al. 2016). For example, lower pleiotropy of DE genes has been found in other cases of rapid adaptation, such as in pupfishes or European grayling (Papakostas et al. 2014; McGirr and Martin 2018). In contrast, our finding that DS genes showed higher pleiotropy and were central in regulatory networks (Fig. 3b,c) suggests that alternative splicing might provide a mechanism through which the function or expression of hub genes and highly pleiotropic genes can be altered or diversified without derailing vital expression patterns or coding sequence (Bush et al. 2017). However, the opposite has been argued in other studies (Iñiguez and Hernández 2017). This discrepancy highlights the need for more large-scale comparative and functional studies of alternative splicing in adaptation to better understand its functional role and the underlying regulatory mechanisms.

### Genetic regulation of alternative splicing and gene expression

Phenotypic divergence between Arctic charr ecotypes has been suggested to be affected by both heritable genetic variation and phenotypic plasticity (Adams and Huntingford 2002b, 2004; Klemetsen 2002). By mapping cis-regulatory expression and splicing QTL, we show that differential expression and differential splicing are at least partially genetically determined, with between 6.7 to 10.1% of DE genes and 16.5% of differentially spliced intron clusters being associated with *cis*-regulatory variation. *Cis*-regulatory divergence has been shown to play an important role in rapid and parallel adaptive divergence in other species (Prud’homme et al. 2007; Wittkopp et al. 2008; Verta and Jones 2019). Consistent with previours findings for cis-regulatory expression QTL in Arctic charr and other postglacial fishes (Rougeux et al. 2019; Verta and Jones 2019; Jacobs et al. 2020), cis-regulatory QTL regulated in multiple cases the differential splicing of genes across replicated ecotypes (Fig. 5b,c), suggesting the importance of shared adaptive regulatory variation in parallel evolution. However, due to our limited sample size, we were only able to map the strongest effect loci and could not detect *trans*-QTL, meaning that our estimate of the genetic regulation of differential regulation is likely an underestimate. In general, the vast majority (94%) of *cis*-regulatory QTL either regulated splicing or expression but not both, consistent with findings in humans (Li et al. 2016). This is likely due to the fact that *cis*-eQTL are mostly located close to transcription start sites, while *cis*-sQTL are often located within gene bodies (Lee and Rio 2015; Li et al. 2016). This difference in regulatory architecture potentially explains why different gene sets were differentially spliced and expressed (Fig. 3a).

Similar to many other studies, we did not find any evidence for signatures of selection within differentially regulated genes (Renaut et al. 2012; Hodgins et al. 2016). Surprisingly, neither *cis*-sQTJ.. nor *cis*-eQTL (Fig. 5a) were significantly associated with signatures of selection. However, we found that sites putatively under selection were most likely located in regulatory regions (e.g. 3’UTR), suggesting that they affect gene expression. One possible explanation is that sites under selection are associated with expression and or splicing divergence in other tissues than muscle, as expression and splicing patterns and regulation often drastically differ across tissues (Saha et al. 2017). Overall, our results suggest that both differential expression and splicing are in part genetically regulated and that these *cis*-regulatory loci are in part parallel across lakes, which is in agreement with our earlier findings (Jacobs et al. 2020).

### Limitations and future work

Our study provides novel insights into the role of post-transcriptional processes in facilitating the rapid and parallel postglacial adaptive divergence in ecological context. Our use of complementary approaches to overcome challenges and limitations of alternative splicing analyses based on short-read data in natural populations of a non-model organism is important because here several lines of evidence support our inferences. While the application of novel analytical approaches to existing data provides new fundamental insights (Jacobs et al. 2018), the rise of long-read data and direct RNA sequencing will facilitate a better understanding of these processes in natural populations.

One important point to be noted is that different isoforms can have different or novel functions that are putatively not annotated (Eksi et al. 2013; Bush et al. 2017). Thus, the GO term based functional inference is limited and likely conservative. Without functional assays of translation and function, which are beyond the scope of this study, we cannot infer the potential of new or different functional roles for different isoforms (Stamm et al. 2005; Mallarino et al. 2017). However, the fact that DS and DE genes were largely associated with different processes and functions suggests that these alternative processes mostly play contrasting roles. Overall, to better understand the role of alternative splicing in rapid ecological adaptation, future work has to focus on i) studying alternative splicing under different adaptive scenarios, ii) identifying the regulatory basis in lab and wild settings and under, ii) using novel long-read approaches to quantify and compare isoform diversity under different evolutionary, genomic and environmental backgrounds, and iv) focus on using functional assays to better understand the downstream impact of alternative splicing.

### Conclusions

Here, we provide important and novel insights into the role of alternative splicing in parallel evolution of natural populations. We suggest that alternative splicing and gene expression in freshwater fish white muscle affect different axes of phenotypic variation; splicing mostly causes structural and functional changes in the muscle and differential expression mostly leading to differences in energy metabolism. Together, we suggest these complementary regulatory processes facilitate the rapid adaptive divergence between Arctic charr ecotypes in foraging and swimming performance. We further show that differentially spliced genes likely play central regulatory roles. This study provides a putative mechanistic framework by which the concerted modification of alternative regulatory processes might facilitate the rapid and parallel adaptive eco-morphological divergence in a postglacial fish.

## Acknowledgments

We thank Madeleine Carruthers, Colin E. Adams and Oliver Hooker for the sample collection that contributed to the original dataset. This work was supported by funding to KRE from Wellcome Trust ISSF (097821/Z/11/Z and 204820/Z/16/Z), Marie Curie CIG 321999 “GEN ECOL ADAPT”, and Carnegie Trust Research Incentive Grant (70287).

## Author contributions

A.J. and K.R.E conceived the project. A.J. conducted all bioinformatic analyses and wrote the initial manuscript. Both authors contributed to the writing of the final manuscript.

## Data accessibility

No new data were generated for this project. The RNA-seq data used for this project are available on NCBI under the BioProject PRJNA551374.

## Notes

### Competing Interest Statement

The authors have declared no competing interest.

