## Supplemental Material for "Alternative splicing and gene expression play contrasting roles in the parallel phenotypic evolution of a salmonid fish"

**Supplementary figures.**

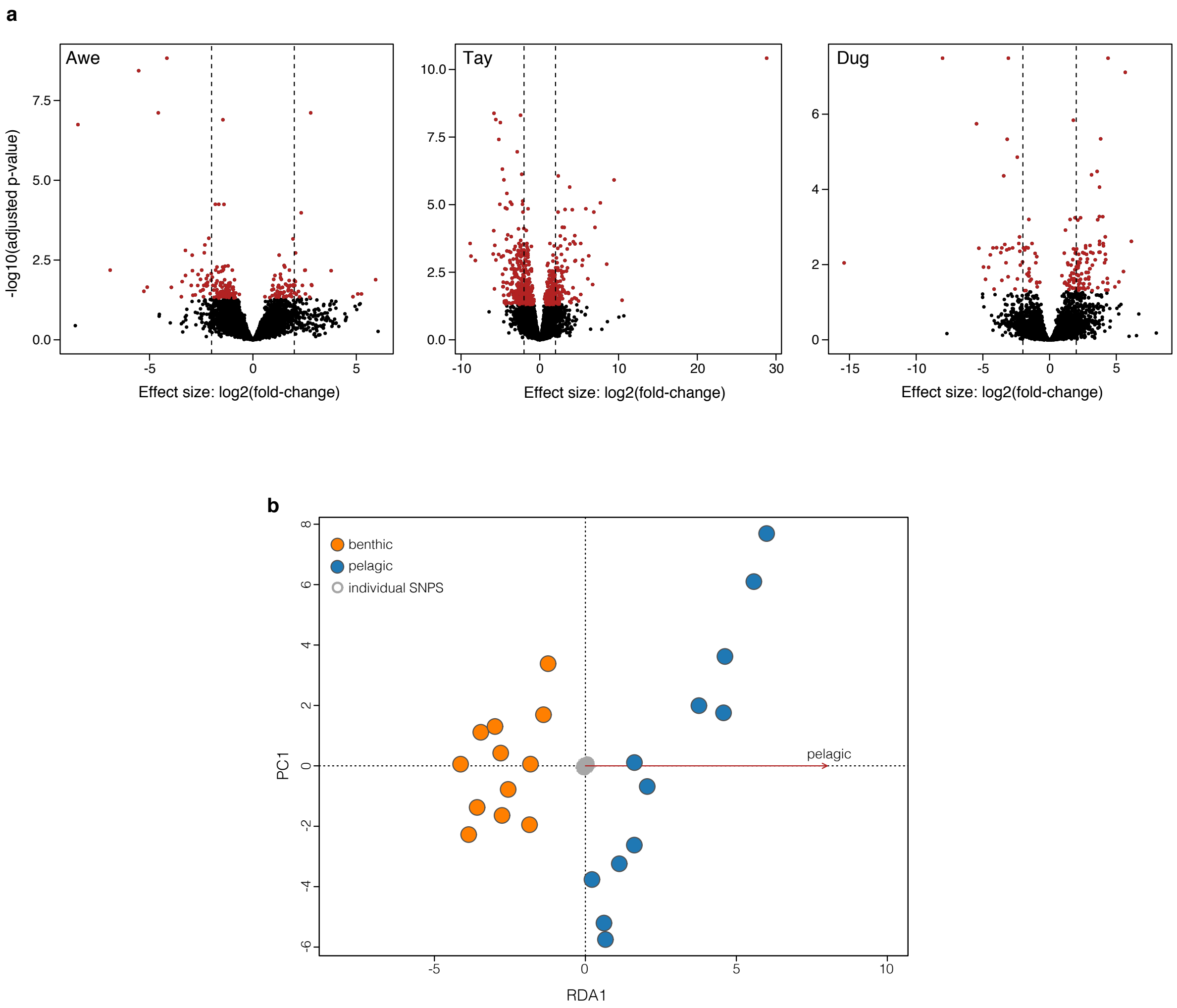


**Fig. S1 - Gene expression divergence.** a) Volcano plots for gene expression divergence between sympatric benthic and pelagic ecotypes in Awe, Tay and Dughaill [Dug]. The plots show the log2(fold-change), representing the effect size and direction of expression divergence, against the adjusted p-value (-log10 transformed). Genes that are overexpressed in the benthic compared to the pelagic ecotype have positive effect sizes. Transcripts that were differentially expressed with a false discovery rate (FDR) below 0.05 are highlighted in red. b) Ordination biplot showing the redundancy axis 1(RDA1) against principal component 1 (PC1). Benthic and pelagic individuals separate along RDA1, while PC1 shows the population structure across lakes.

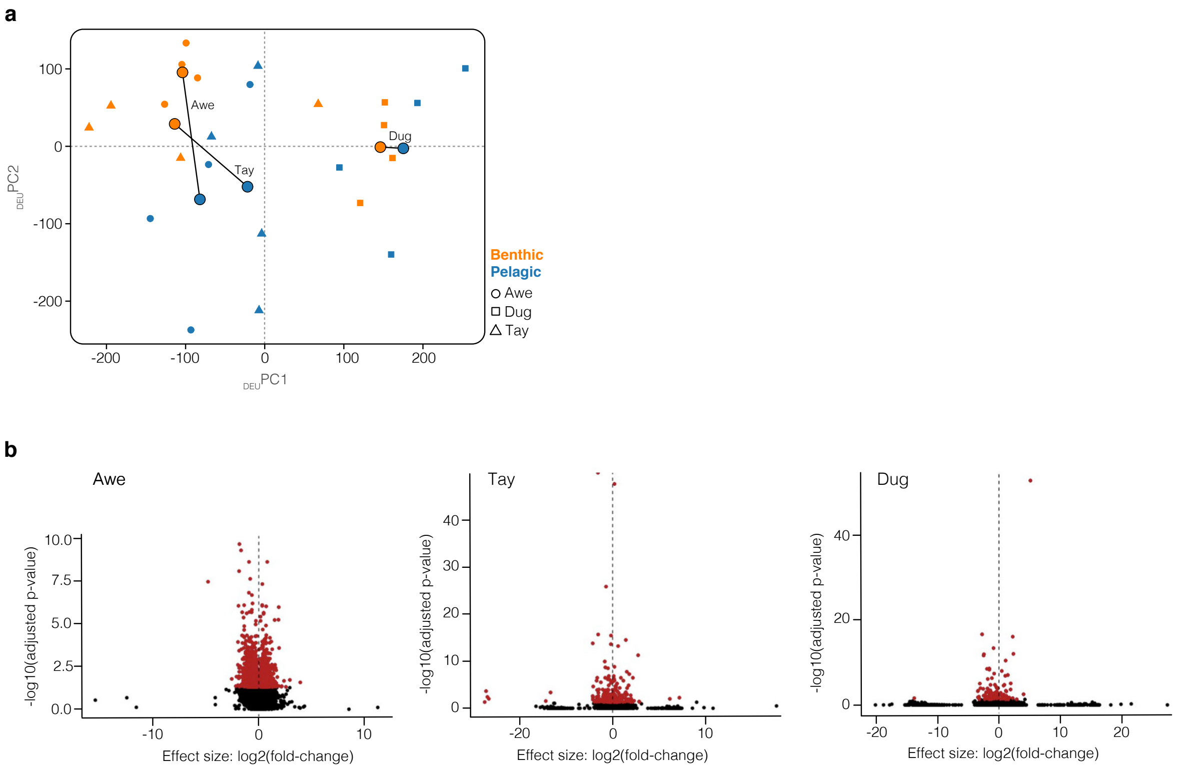


**Fig. S2 - Alternative splicing. a**) Principal components plot for PC1 and PC2 based on rlog-normalised exon-based transcript count data (DEXseq results). Large dots show the centroid for each ecotype (distinguished by colour) by lake, with sympatric ecotypes connected by lines. Small points show individual data points with individuals from different lakes being distinguished by different shapes. **b**) Volcano plots for exon expression data for each lake. See Fig. S1b for plot descriptions. Exons showing significant differences (FDR < 0.05) in exon usage between sympatric ecotypes are highlighted in red.

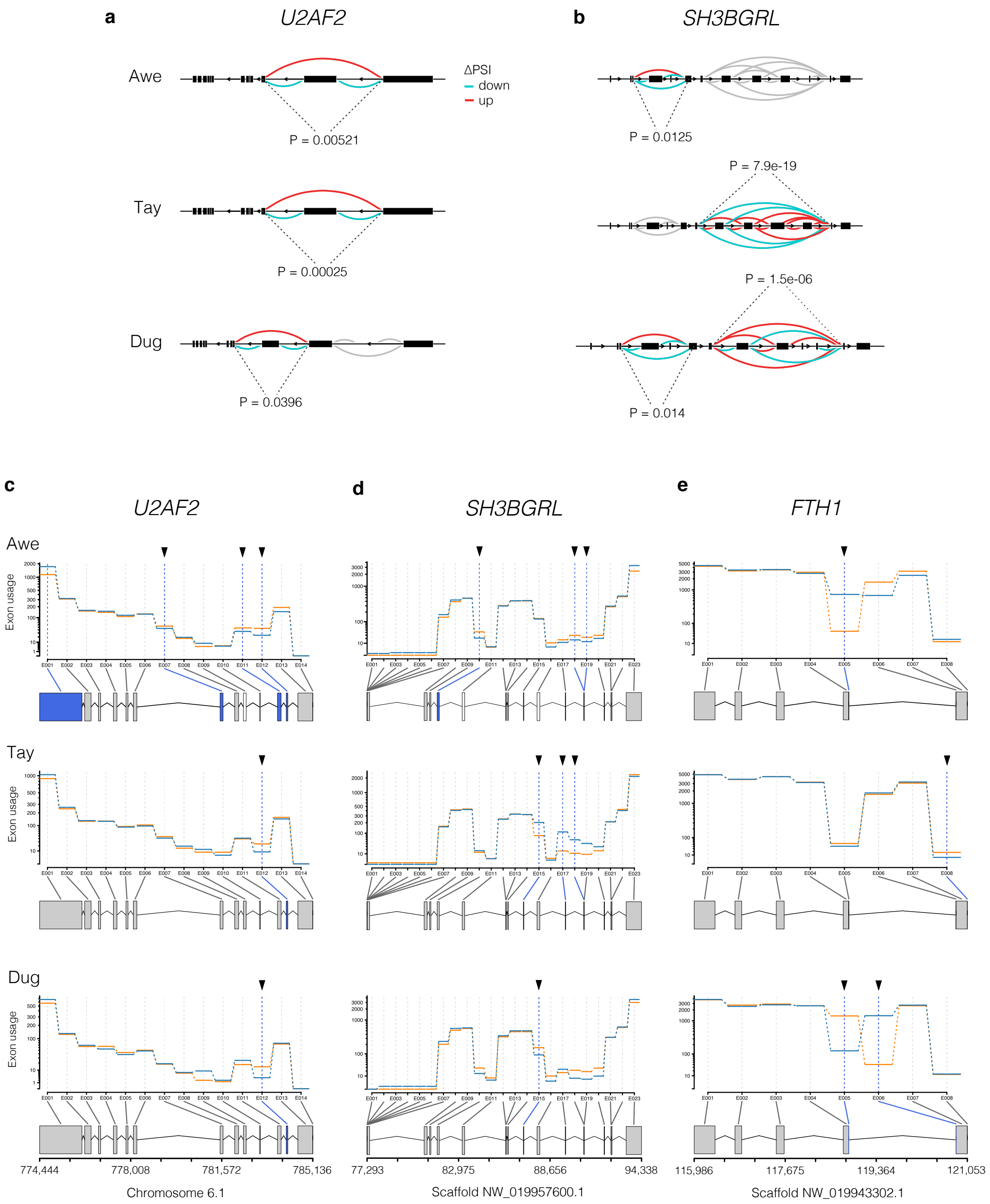


**Fig. S3 - Genes models for shared alternatively spliced genes. a,b,c**) Gene models illustrating alternative splicing patterns for **a**) *U2AF2* and **b**) *SH3BGRL* and **c**) *FTH1* in Awe, Tay and Dug, respectively. Exon usage (y-axis), the expression of each exon corrected for overall gene expression, is shown for each ecotype (orange line for benthic and blue line for pelagic ecotypes) and exons (x-axis) that are differentially spliced are marked with triangles and highlighted in blue in the gene model. **d,e**) Sashimi graphs highlighting patterns of differential intron excision between sympatric ecotypes by lake for intron clusters in **e**) *U2AF2* and **f**) *SH3BGRL*. The amount of differential splicing is measured as 'change in the percent spliced in (ΔPSI)'. For example, in *U2AF2*, one intron cluster shows the increased skipping of one exon (decreased usage) in the benthic ecotype compared to the pelagic ecotype. Although the differentially spliced intron clusters are the same in Awe and Tay, a different intron cluster is differentially spliced in Dug. Associated p-values are shown in the graph.

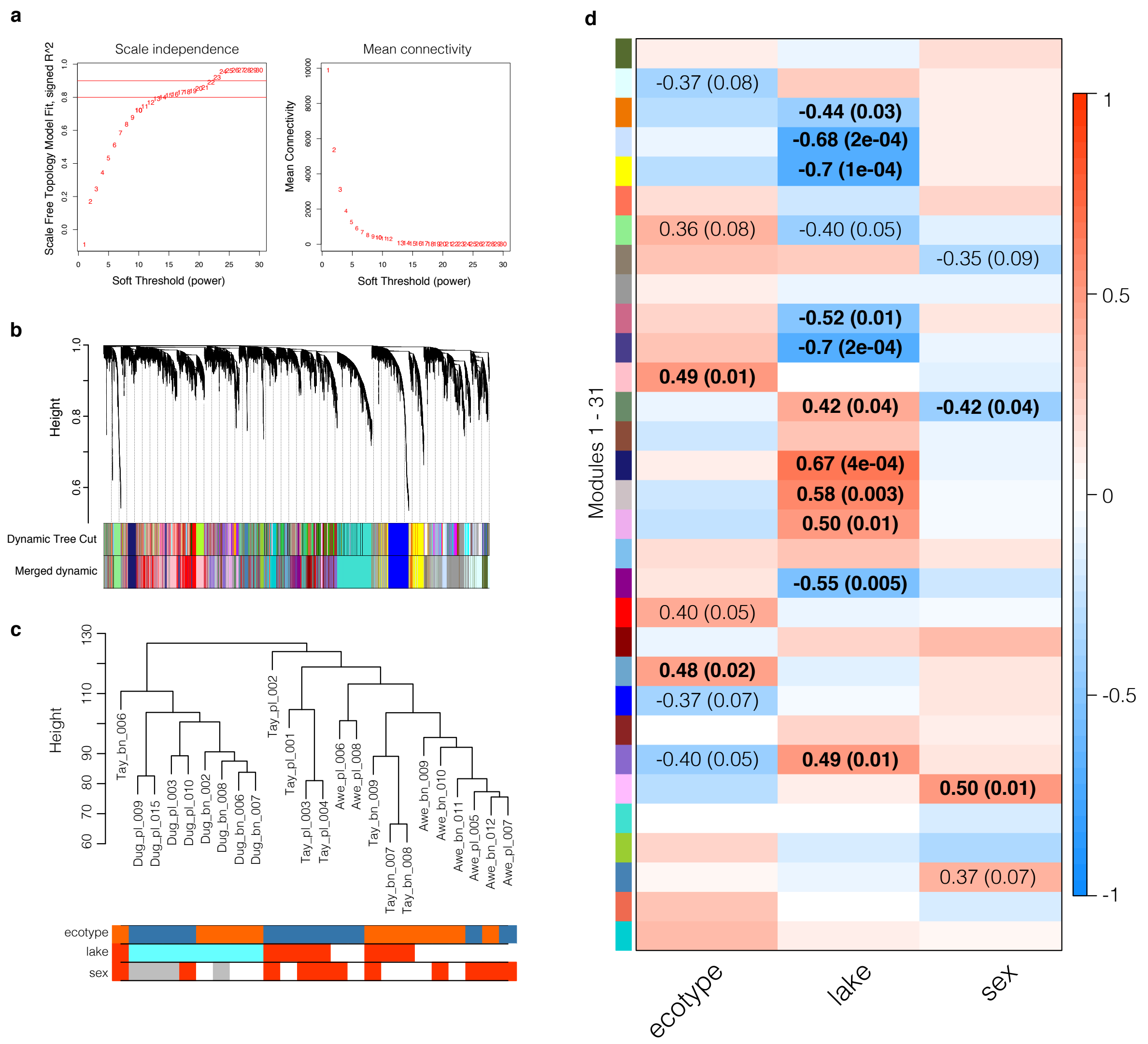


**Fig. S4 - WGCNA gene co-expression results. a**) Choice of soft power for scale free network construction. The soft power was chosen according to the biologically motivated scale free topology criterion, selecting the lowest integer above the model fit $R^2$=0.9. **b**) Hierarchical clustering of genes and identification of eigengene modules using the dynamic tree cut algorithm. Dendrogram based on the topological overlap distance in gene expression profiles for each gene, with branches corresponding to individual identified modules. Identified 'raw' modules (top row in colour bar) and merged modules (bottom row in colour bar) with a similarity above 0.8 are highlighted by the different colours. **c**) Dendrogram based on gene expression similarity between individual samples. Individuals mostly clusters by lake and ecotype. The colour bars show the ecotype, lake of origin and sex for each individual. Individuals with unknown sex are highlighted in grey. **d**) Correlations between module eigengenes and ecotype, lake and sex. Each cell corresponds to the Pearson correlation between a module (shown by colour on the left) and each tested variable, with cells being colour-coded based on the correlation coefficient (scale on the right). Correlation coefficients and p-values (in brackets) are shown for cells with p-values below 0.1 and highlighted in bold for significant correlations ($P < 0.05$).

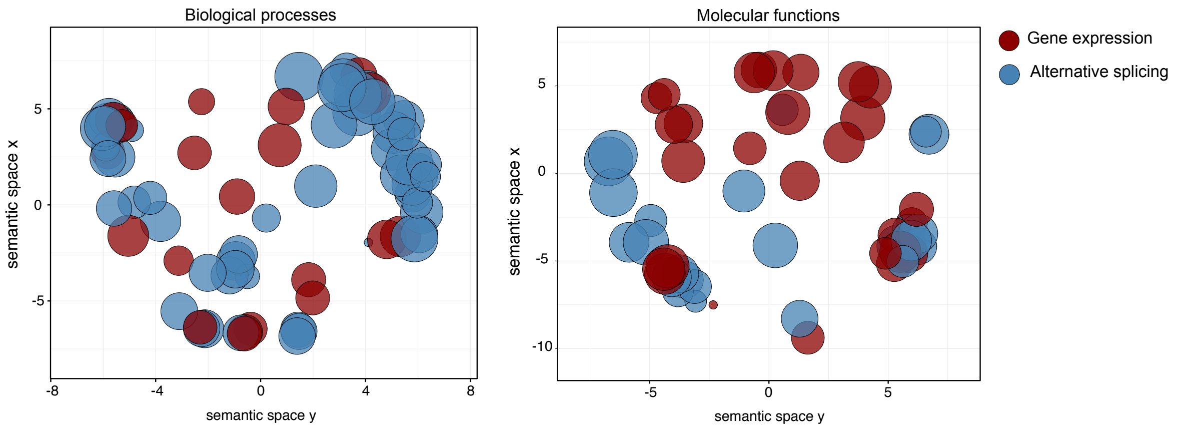


**Fig. S5 - Comparison of enriched gene ontology terms.** Multidimensional scaling plots based on semantic similarities for gene ontology (GO) terms enriched for alternatively spliced genes (blue circles; based on leafcutter and DEXseq) and differentially expressed genes (red circles). The plots highlight the functional differences in biological processes (left) and molecular functions (right) for genes that are differentially spliced or differentially expressed between benthic and pelagic ecotypes. Each circle represents one gene ontology term with the size of the circle corresponding to the size of the gene ontology term (number of genes associated with a GO term).

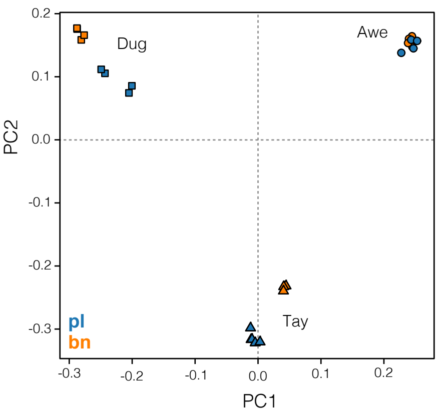


**Fig. S6 – Principal components analysis based on LD-pruned SNPs.** Biplot showing PC1 against PC2 based on a LD-pruned transcriptome-based SNP dataset (n = 12,864 SNPs) for all individuals. Ecotypes are coded by colour and lakes are coded by shape.

### Supplementary Tables:

**Table S1 – Ecotype-associated differentially expressed genes.** List of genes with ecotype-associated expression patterns that are also differentially expressed in two populations.

| Gene ID | LFC-Awe | LFC-Tay | LFC-Dug | RDA z-score | Gene symbol |
|---|---|---|---|---|---|
| rna1201 | -5.11 | -5.76 | -1.82 | 2.24 | *HOXC11* |
| rna13255 | -1.74 | -0.77 | -1.55 | 2.82 | *EIF2S2* |
| rna13260 | -1.45 | -0.05 | -0.96 | 2.77 | *SRSF6* |
| rna13908 | 0.92 | 0.89 | 0.94 | -2.83 | *PPP4R2* |
| rna15895 | 1.01 | 4.45 | 3.78 | -2.31 | *PVALB* |
| rna16690 | -0.08 | -1.33 | -2.42 | 2.35 | *BEND3* |
| rna2035 | -1.00 | -4.17 | -4.80 | 2.42 | *HESX1* |
| rna22158 | 1.17 | 1.49 | -0.02 | -2.22 | *ZBTB2* |
| rna22829 | 3.78 | 2.61 | 4.04 | -2.39 | *EGFR* |
| rna23988 | -2.20 | -2.69 | 0.26 | 2.18 | *FGF17* |
| rna30070 | 2.79 | 1.68 | -0.12 | -2.25 | *PPRC1* |
| rna36918 | -1.46 | -1.51 | -0.62 | 2.86 | *RNF7* |
| rna39482 | 0.94 | 0.82 | 0.12 | -2.04 | *TP53* |
| rna45229 | 1.75 | 0.49 | 2.32 | -2.79 | *JARID2* |
| rna51935 | 1.43 | 1.57 | 0.32 | -2.37 | *PTPN4* |
| rna52559 | 0.26 | 4.13 | 2.66 | -2.02 | *EPD* |
| rna55602 | 2.06 | 1.86 | 0.78 | -2.87 | *ZNF-C2H2-type* |
| rna56374 | 0.07 | 3.81 | 4.37 | -2.34 | *MYH* |
| rna57430 | -1.99 | -3.70 | -3.21 | 2.70 | *COL11A1* |
| rna59180 | -1.00 | -0.82 | -0.31 | 2.62 | *PON* |
| rna62721 | -0.34 | -1.99 | -3.36 | 2.61 | *SOD1* |
| rna772 | -0.33 | -1.21 | -1.32 | 2.66 | *APOBEC2* |
| rna8525 | -2.25 | -1.95 | -0.63 | 2.82 | *GALNT8* |

Notes: LFC = Log fold change in Awe, Tay and Dug. Genes with negative LFC and a positive RDA z-score show higher expression in the pelagic ecotype and genes with positive LFC and negative z-score show higher expression in the benthic ecotype.

**Table S2 – Shared differentially spliced genes.** List of genes that were identified as differentially spliced in at least two populations with at least one of the two methods.

| Transcript | Populations | Method | Gene symbol |
|---|---|---|---|
| rna11110 | All | DEU + DIE | *U2AF2* |
| rna67268 | All | DEU + DIE | *SH3BGRL* |
| rna58592 | All | DEU | *FTH1* |
| rna12079 | Awe + Dug | DIE | *CSNK1A1* |
| rna13739 | Awe + Dug | DIE | *AMPD1* |
| rna13742 | Awe + Dug | DIE | *CSDE1* |
| rna19299 | Awe + Dug | DIE | *ADIPOR1* |
| rna21198 | Awe + Dug | DIE | *SERBP1* |
| rna22297 | Awe + Dug | DIE | *KINX* |
| rna22935 | Awe + Dug | DIE | *RPL21* |
| rna30130 | Awe + Dug | DIE | *EIF3A* |
| rna30306 | Awe + Dug | DIE | *MYOZ1* |
| rna30655 | Awe + Dug | DIE | *RTN4* |
| rna32110 | Awe + Dug | DIE | *MARCH6* |
| rna36381 | Awe + Dug | DIE | *EIF4H* |
| rna38208 | Awe + Dug | DIE | *USP28* |
| rna40224 | Awe + Dug | DIE | *EIF3A* |
| rna41389 | Awe + Dug | DIE | *CIRBPB* |
| rna27736 | Awe + Tay | DIE | *NACA* |
| rna36761 | Awe + Tay | DIE | *USP2* |
| rna54911 | Awe + Tay | DIE | *GSPT1* |
| rna56373 | Awe + Tay | DIE | *MYH* |
| rna2022 | Tay + Dug | DIE | *IGFN1* |
| rna24607 | Tay + Dug | DIE | *LOC111974326* |
| rna30709 | Tay + Dug | DIE | *RPS24* |
| rna37452 | Tay + Dug | DIE | *SH3BGRL2* |
| rna40780 | Tay + Dug | DIE | *RTN4* |
| rna44131 | Tay + Dug | DIE | *TRDNL* |
| rna5821 | Tay + Dug | DIE | *TMEM38B* |

| | | | |
|---|---|---|---|
| rna67221 | Tay + Dug | DIE | *SRSF7* |
| rna10966 | Awe + Dug | DEU | *RBM22* |
| rna13937 | Awe + Dug | DEU | *LAP2B* |
| rna14329 | Awe + Dug | DEU | *NACA* |
| rna27088 | Awe + Dug | DEU | *PYCR3L* |
| rna32442 | Awe + Dug | DEU | *AGL* |
| rna34376 | Awe + Dug | DEU | *PIGL* |
| rna43091 | Awe + Dug | DEU | *MYOM1* |
| rna45075 | Awe + Dug | DEU | *PPP1R10* |
| rna1016 | Awe + Tay | DEU | *TRIB3* |
| rna17669 | Awe + Tay | DEU | *MTSS1L* |
| rna18421 | Awe + Tay | DEU | *RBM5* |
| rna18723 | Awe + Tay | DEU | *PFKFB4* |
| rna19665 | Awe + Tay | DEU | *ICAL* |
| rna2033 | Awe + Tay | DEU | *APPL1* |
| rna21099 | Awe + Tay | DEU | *PALLD* |
| rna22238 | Awe + Tay | DEU | *LAMA2* |
| rna2628 | Awe + Tay | DEU | *CNP* |
| rna26502 | Awe + Tay | DEU | *GNA11* |
| rna31887 | Awe + Tay | DEU | *OBSL* |
| rna3224 | Awe + Tay | DEU | *BCORL* |
| rna364 | Awe + Tay | DEU | *ATL2* |
| rna36842 | Awe + Tay | DEU | *NUMBL* |
| rna40182 | Awe + Tay | DEU | *HERC4* |
| rna44298 | Awe + Tay | DEU | *CAD* |
| rna48617 | Awe + Tay | DEU | *UBAP2* |
| rna4939 | Awe + Tay | DEU | *C4L* |
| rna554 | Awe + Tay | DEU | *TXNRD1* |
| rna57514 | Awe + Tay | DEU | *FLNCL* |
| rna60467 | Awe + Tay | DEU | *CNOT3* |
| rna62138 | Awe + Tay | DEU | *RBM20* |
| rna66230 | Awe + Tay | DEU | *MYH* |

| | | | |
|---|---|---|---|
| rna7128 | Awe + Tay | DEU | *PRRC1* |
| rna7911 | Awe + Tay | DEU | *WWP2* |
| rna61622 | Tay + Dug | DEU | *LOC112075266* |
| rna14241 | Awe + Dug | DEU + DIE | *HNRNPA1* |
| rna2334 | Awe + Dug | DEU + DIE | *LOC111974535* |
| rna28599 | Awe + Dug | DEU + DIE | *HNRNPA1* |
| rna32378 | Awe + Dug | DEU + DIE | *ASPH* |
| rna37159 | Awe + Dug | DEU + DIE | *DCUN1D5* |
| rna52530 | Awe + Dug | DEU + DIE | *YBX2A* |
| rna51879 | Tay + Dug | DEU + DIE | *NEB* |

Notes: Methods - Differential Exon Usage (DEU) in *DEXseq*, Differential Intron Excision (DIE) in *leafcutter*. Genes without gene symbol show the locus ID from the Arctic charr genome annotation.

**Table S3 – Top and bottom 20 enriched GO terms for genes with ecotype-associated expression patterns (identified in the redundancy analysis).**

| GO ID | GO description | Size | NES | FDR |
|---|---|---|---|---|
| GO:0006119 | oxidative phosphorylation | 193 | 6.68 | <0.0001 |
| GO:0042773 | ATP synthesis coupled electron transport | 124 | 6.51 | <0.0001 |
| GO:0002181 | cytoplasmic translation | 177 | 6.38 | <0.0001 |
| GO:0042775 | mitochondrial ATP synthesis coupled electron transport | 124 | 6.30 | <0.0001 |
| GO:0022904 | respiratory electron transport chain | 158 | 6.27 | <0.0001 |
| GO:0006414 | translational elongation | 156 | 6.13 | <0.0001 |
| GO:0033108 | mitochondrial respiratory chain complex assembly | 112 | 6.09 | <0.0001 |
| GO:0022900 | electron transport chain | 230 | 6.09 | <0.0001 |
| GO:0006120 | mitochondrial electron transport, NADH to ubiquinone | 65 | 5.98 | <0.0001 |
| GO:0010257 | NADH dehydrogenase complex assembly | 72 | 5.52 | <0.0001 |
| GO:0046034 | ATP metabolic process | 443 | 5.46 | <0.0001 |
| GO:0032981 | mitochondrial respiratory chain complex I assembly | 72 | 5.43 | <0.0001 |
| GO:0009126 | purine nucleoside monophosphate metabolic process | 496 | 5.28 | <0.0001 |
| GO:0009123 | nucleoside monophosphate metabolic process | 521 | 5.22 | <0.0001 |
| GO:0009167 | purine ribonucleoside monophosphate metabolic process | 494 | 5.20 | <0.0001 |
| GO:0009161 | ribonucleoside monophosphate metabolic process | 506 | 5.07 | <0.0001 |
| GO:0070125 | mitochondrial translational elongation | 89 | 5.06 | <0.0001 |
| GO:0070126 | mitochondrial translational termination | 90 | 5.02 | <0.0001 |
| GO:0009141 | nucleoside triphosphate metabolic process | 557 | 5.00 | <0.0001 |
| GO:0032543 | mitochondrial translation | 140 | 4.98 | <0.0001 |
| GO:0043406 | positive regulation of MAP kinase activity | 469 | -3.30 | <0.0001 |
| GO:0048569 | post-embryonic animal organ development | 331 | -3.31 | <0.0001 |
| GO:0034332 | adherens junction organization | 318 | -3.31 | <0.0001 |
| GO:0035264 | multicellular organism growth | 404 | -3.31 | <0.0001 |
| GO:0003158 | endothelium development | 343 | -3.32 | <0.0001 |
| GO:0046643 | regulation of gamma-delta T cell activation | 61 | -3.35 | <0.0001 |
| GO:0050701 | interleukin-1 secretion | 121 | -3.36 | <0.0001 |
| GO:0048737 | imaginal disc-derived appendage development | 169 | -3.38 | <0.0001 |

| | | | | |
|---|---|---|---|---|
| GO:0045586 | regulation of gamma-delta T cell differentiation | 61 | -3.38 | <0.0001 |
| GO:0002699 | positive regulation of immune effector process | 340 | -3.41 | <0.0001 |
| GO:0046578 | regulation of Ras protein signal transduction | 357 | -3.42 | <0.0001 |
| GO:0032147 | activation of protein kinase activity | 489 | -3.48 | <0.0001 |
| GO:0050707 | regulation of cytokine secretion | 305 | -3.48 | <0.0001 |
| GO:0046777 | protein autophosphorylation | 471 | -3.57 | <0.0001 |
| GO:0045619 | regulation of lymphocyte differentiation | 404 | -3.74 | <0.0001 |
| GO:0043547 | positive regulation of GTPase activity | 489 | -3.76 | <0.0001 |
| GO:0007160 | cell-matrix adhesion | 426 | -3.80 | <0.0001 |
| GO:0051056 | regulation of small GTPase mediated signal transduction | 504 | -3.82 | <0.0001 |
| GO:0050663 | cytokine secretion | 368 | -3.86 | <0.0001 |
| GO:0045807 | positive regulation of endocytosis | 351 | -3.86 | <0.0001 |

Note: NES = Normalized enrichment score. FDR = False discovery rate.

**Table S4 – GO terms overrepresented for ecotype-associated genes.** None of the GO terms were significantly overrepresented after correction for multiple testing (FDR < 0.05).

| GO ID | GO description | log10(p-value) | uniqueness |
|---|---|---|---|
| GO:0008152 | metabolic process | -2.31 | 1.00 |
| GO:0009987 | cellular process | -1.73 | 1.00 |
| GO:0044238 | primary metabolic process | -1.78 | 0.98 |
| GO:0071704 | organic substance metabolic process | -1.79 | 0.98 |
| GO:1900673 | olefin metabolic process | -3.68 | 0.98 |
| GO:0035526 | retrograde transport, plasma membrane to Golgi | -1.88 | 0.96 |
| GO:1901360 | organic cyclic compound metabolic process | -3.25 | 0.95 |
| GO:0070988 | demethylation | -2.19 | 0.95 |
| GO:1900619 | acetate ester metabolic process | -1.84 | 0.94 |
| GO:0044237 | cellular metabolic process | -2.85 | 0.94 |
| GO:0042982 | amyloid precursor protein metabolic process | -1.65 | 0.94 |
| GO:0006556 | S-adenosylmethionine biosynthetic process | -1.58 | 0.93 |
| GO:0018904 | ether metabolic process | -1.48 | 0.93 |
| GO:0009693 | ethylene biosynthetic process | -2.38 | 0.93 |
| GO:0002175 | protein localization to paranode region of axon | -2.04 | 0.92 |
| GO:0097332 | response to antipsychotic drug | -1.68 | 0.92 |
| GO:0046483 | heterocycle metabolic process | -3.32 | 0.92 |
| GO:0006725 | cellular aromatic compound metabolic process | -2.98 | 0.92 |
| GO:0050435 | beta-amyloid metabolic process | -2.83 | 0.92 |
| GO:0010467 | gene expression | -1.52 | 0.92 |
| GO:0070269 | pyroptosis | -2.85 | 0.91 |
| GO:0051725 | protein de-ADP-ribosylation | -2.38 | 0.91 |
| GO:0010288 | response to lead ion | -1.64 | 0.91 |
| GO:0009411 | response to UV | -2.34 | 0.91 |
| GO:0006713 | glucocorticoid catabolic process | -2.85 | 0.91 |
| GO:0009719 | response to endogenous stimulus | -2.56 | 0.91 |
| GO:0009404 | toxin metabolic process | -2.50 | 0.91 |
| GO:0032196 | transposition | -1.60 | 0.91 |
| GO:0097176 | epoxide metabolic process | -1.71 | 0.90 |

| | | | |
|---|---|---|---|
| GO:0006623 | protein targeting to vacuole | -2.66 | 0.90 |
| GO:0043696 | dedifferentiation | -1.65 | 0.90 |
| GO:0031365 | N-terminal protein amino acid modification | -2.50 | 0.89 |
| GO:0045472 | response to ether | -2.38 | 0.89 |
| GO:0045116 | protein neddylation | -4.55 | 0.89 |
| GO:0034552 | respiratory chain complex II assembly | -2.09 | 0.89 |
| GO:0006474 | N-terminal protein amino acid acetylation | -2.84 | 0.89 |
| GO:0072527 | pyrimidine-containing compound metabolic process | -1.68 | 0.89 |
| GO:0009087 | methionine catabolic process | -1.58 | 0.88 |
| GO:0034641 | cellular nitrogen compound metabolic process | -4.26 | 0.88 |
| GO:0009193 | pyrimidine ribonucleoside diphosphate metabolic process | -1.47 | 0.88 |
| GO:0030205 | dermatan sulfate metabolic process | -1.48 | 0.88 |
| GO:0006369 | termination of RNA polymerase II transcription | -2.21 | 0.88 |
| GO:0036058 | filtration diaphragm assembly | -1.88 | 0.87 |
| GO:0043933 | macromolecular complex subunit organization | -1.83 | 0.87 |
| GO:0070647 | protein modification by small protein conjugation or removal | -1.59 | 0.87 |
| GO:0032776 | DNA methylation on cytosine | -2.22 | 0.87 |
| GO:0043697 | cell dedifferentiation | -1.65 | 0.87 |
| GO:0070966 | nuclear-transcribed mRNA catabolic process, no-go decay | -3.30 | 0.87 |
| GO:0009838 | abscission | -1.54 | 0.86 |
| GO:0072528 | pyrimidine-containing compound biosynthetic process | -1.70 | 0.86 |
| GO:0090305 | nucleic acid phosphodiester bond hydrolysis | -1.50 | 0.86 |
| GO:0006353 | DNA-templated transcription, termination | -1.72 | 0.86 |
| GO:0043388 | positive regulation of DNA binding | -3.20 | 0.85 |
| GO:0008380 | RNA splicing | -3.46 | 0.85 |
| GO:0006259 | DNA metabolic process | -2.73 | 0.84 |
| GO:0010506 | regulation of autophagy | -1.52 | 0.84 |
| GO:1905563 | negative regulation of vascular endothelial cell proliferation | -1.58 | 0.84 |
| GO:1901699 | cellular response to nitrogen compound | -3.22 | 0.84 |
| GO:0007422 | peripheral nervous system development | -3.20 | 0.84 |

| | | | |
|---|---|---|---|
| GO:0010528 | regulation of transposition | -1.81 | 0.83 |
| GO:0006396 | RNA processing | -1.63 | 0.83 |
| GO:0000454 | snoRNA guided rRNA pseudouridine synthesis | -1.88 | 0.83 |
| GO:0042445 | hormone metabolic process | -2.74 | 0.83 |
| GO:1902996 | regulation of neurofibrillary tangle assembly | -2.85 | 0.81 |
| GO:0060408 | regulation of acetylcholine metabolic process | -2.09 | 0.80 |
| GO:2000325 | regulation of ligand-dependent nuclear receptor transcription coactivator activity | -1.88 | 0.80 |
| GO:1902954 | regulation of early endosome to recycling endosome transport | -2.85 | 0.80 |
| GO:0044878 | mitotic cytokinesis checkpoint | -1.94 | 0.80 |
| GO:0040029 | regulation of gene expression, epigenetic | -2.97 | 0.80 |
| GO:0010197 | polar nucleus fusion | -2.09 | 0.79 |
| GO:1902947 | regulation of tau-protein kinase activity | -3.66 | 0.79 |
| GO:0034371 | chylomicron remodeling | -2.38 | 0.77 |
| GO:1902187 | negative regulation of viral release from host cell | -3.78 | 0.77 |
| GO:1902268 | negative regulation of polyamine transmembrane transport | -1.88 | 0.77 |
| GO:0090342 | regulation of cell aging | -1.83 | 0.77 |
| GO:1901898 | negative regulation of relaxation of cardiac muscle | -2.42 | 0.76 |
| GO:0019227 | neuronal action potential propagation | -2.07 | 0.76 |
| GO:2001020 | regulation of response to DNA damage stimulus | -3.71 | 0.75 |
| GO:0010823 | negative regulation of mitochondrion organization | -2.90 | 0.74 |
| GO:1902430 | negative regulation of beta-amyloid formation | -3.78 | 0.74 |
| GO:0045935 | positive regulation of nucleobase-containing compound metabolic process | -2.99 | 0.68 |

Note: log10(p-value) = log-transformed p-value; uniqueness = GO terms with a higher uniqueness are less common in the overall dataset. Calculated as 1-[average semantic similarity with all other GO terms].

**Table S5 – Shared GO terms overrepresented for differentially spliced genes.**

| GO ID | GO description | GO category |
|---|---|---|
| GO:0006936 | muscle contraction | biological processes |
| GO:0050658 | RNA transport | biological processes |
| GO:0061061 | muscle structure development | biological processes |
| GO:1903311 | regulation of mRNA metabolic process | biological processes |
| GO:0031032 | actomyosin structure organization | biological processes |
| GO:0006413 | translational initiation | biological processes |
| GO:1905580 | positive regulation of ERBB3 signaling pathway | biological processes |
| GO:0019439 | aromatic compound catabolic process | biological processes |
| GO:0030029 | actin filament-based process | biological processes |
| GO:0010467 | gene expression | biological processes |
| GO:0043603 | cellular amide metabolic process | biological processes |
| GO:0071243 | cellular response to arsenic-containing substance | biological processes |
| GO:0008380 | RNA splicing | biological processes |
| GO:1901576 | organic substance biosynthetic process | biological processes |
| GO:0097010 | eukaryotic translation initiation factor 4F complex assembly | biological processes |
| GO:0016071 | mRNA metabolic process | biological processes |
| GO:0010608 | posttranscriptional regulation of gene expression | biological processes |
| GO:0090158 | endoplasmic reticulum membrane organization | biological processes |
| GO:0097435 | supramolecular fiber organization | biological processes |
| GO:0006396 | RNA processing | biological processes |
| GO:1903608 | protein localization to cytoplasmic stress granule | biological processes |
| GO:0002183 | cytoplasmic translational initiation | biological processes |
| GO:0051168 | nuclear export | biological processes |
| GO:0002181 | cytoplasmic translation | biological processes |
| GO:0051254 | positive regulation of RNA metabolic process | biological processes |
| GO:0034248 | regulation of cellular amide metabolic process | biological processes |
| GO:0006403 | RNA localization | biological processes |
| GO:0010927 | cellular component assembly involved in morphogenesis | biological processes |
| GO:0014706 | striated muscle tissue development | biological processes |
| GO:0006417 | regulation of translation | biological processes |

| | | |
|---|---|---|
| GO:0010468 | regulation of gene expression | biological processes |
| GO:0042692 | muscle cell differentiation | biological processes |
| GO:0003007 | heart morphogenesis | biological processes |
| GO:0022618 | ribonucleoprotein complex assembly | biological processes |
| GO:0034645 | cellular macromolecule biosynthetic process | biological processes |
| GO:0009059 | macromolecule biosynthetic process | biological processes |
| GO:1903312 | negative regulation of mRNA metabolic process | biological processes |
| GO:0031329 | regulation of cellular catabolic process | biological processes |
| GO:0044271 | cellular nitrogen compound biosynthetic process | biological processes |
| GO:0000380 | alternative mRNA splicing, via spliceosome | biological processes |
| GO:0015931 | nucleobase-containing compound transport | biological processes |
| GO:0019222 | regulation of metabolic process | biological processes |
| GO:0031034 | myosin filament assembly | biological processes |
| GO:0010605 | negative regulation of macromolecule metabolic process | biological processes |
| GO:0031033 | myosin filament organization | biological processes |
| GO:0007029 | endoplasmic reticulum organization | biological processes |
| GO:0022607 | cellular component assembly | biological processes |
| GO:0071826 | ribonucleoprotein complex subunit organization | biological processes |
| GO:0043933 | macromolecular complex subunit organization | biological processes |
| GO:0060047 | heart contraction | biological processes |
| GO:0043484 | regulation of RNA splicing | biological processes |
| GO:0070925 | organelle assembly | biological processes |
| GO:0006412 | translation | biological processes |
| GO:0033120 | positive regulation of RNA splicing | biological processes |
| GO:0003012 | muscle system process | biological processes |
| GO:0030048 | actin filament-based movement | biological processes |
| GO:0005198 | structural molecule activity | molecular function |
| GO:0008307 | structural constituent of muscle | molecular function |
| GO:0042805 | actinin binding | molecular function |
| GO:0061752 | telomeric repeat-containing RNA binding | molecular function |
| GO:0043021 | ribonucleoprotein complex binding | molecular function |
| GO:0060090 | binding, bridging | molecular function |

| | | |
|---|---|---|
| GO:0044877 | macromolecular complex binding | molecular function |
| GO:0097159 | organic cyclic compound binding | molecular function |
| GO:0097617 | annealing activity | molecular function |
| GO:0003697 | single-stranded DNA binding | molecular function |
| GO:0003676 | nucleic acid binding | molecular function |
| GO:0003723 | RNA binding | molecular function |
| GO:0003727 | single-stranded RNA binding | molecular function |
| GO:1901363 | heterocyclic compound binding | molecular function |
| GO:0033592 | RNA strand annealing activity | molecular function |
| GO:0036002 | pre-mRNA binding | molecular function |
| GO:0008092 | cytoskeletal protein binding | molecular function |
| GO:0097718 | disordered domain specific binding | molecular function |
| GO:0003743 | translation initiation factor activity | molecular function |
| GO:0003725 | double-stranded RNA binding | molecular function |
| GO:0003729 | mRNA binding | molecular function |
| GO:0030674 | protein binding, bridging | molecular function |
| GO:0051371 | muscle alpha-actinin binding | molecular function |
| GO:0019904 | protein domain specific binding | molecular function |
| GO:0008135 | translation factor activity, RNA binding | molecular function |
| GO:0043024 | ribosomal small subunit binding | molecular function |
| GO:0003779 | actin binding | molecular function |

**Table S6 - *Cis*-sQTL for differentially spliced genes in at least two populations.** Only the SNP with the strongest association with intron excision ratio is shown.

| intron cluster | SNP | F-statistic | FDR | beta | $R^2$ | Transcript |
|---|---|---|---|---|---|---|
| clu_1082_NA | rs9218 | -6.45 | 2.6E-03 | -1.37 | 0.66 | rna19299 |
| clu_1127_NA | rs9326 | -4.75 | 4.2E-02 | -1.67 | 0.52 | rna19665 |
| clu_1176_NA | rs32035 | 5.52 | 1.2E-02 | 1.19 | 0.59 | rna59180 |
| clu_1359_NA | rs42959 | 6.24 | 3.6E-03 | 1.10 | 0.65 | rna67221 |
| clu_1557_NA | rs13626 | -4.71 | 4.4E-02 | -2.06 | 0.51 | rna29511 |
| clu_1562_NA | rs13658 | -5.43 | 1.4E-02 | -0.98 | 0.58 | rna29596 |
| clu_1642_NA | rs13843 | 5.89 | 6.3E-03 | 1.09 | 0.62 | rna30305 |
| clu_1758_NA | rs30784 | -5.07 | 2.4E-02 | -1.29 | 0.55 | NA |
| clu_1965_NA | rs12977 | -5.65 | 9.5E-03 | -0.92 | 0.60 | rna27735 |
| clu_2200_NA | rs35027 | 5.38 | 1.5E-02 | 3.40 | 0.58 | rna61622 |
| clu_244_NA | rs11638 | 7.06 | 9.1E-04 | 3.07 | 0.70 | rna24559 |
| clu_2595_NA | rs8640 | -12.45 | 4.3E-07 | -1.06 | 0.88 | NA |
| clu_2614_NA | rs8760 | 26.54 | 4.0E-12 | 1.14 | 0.97 | rna18120 |
| clu_2692_NA | rs38691 | 6.61 | 1.9E-03 | 1.69 | 0.68 | rna63801 |
| clu_2864_NA | rs6546 | 4.77 | 4.0E-02 | 1.07 | 0.52 | rna13561 |
| clu_300_NA | rs11817 | 4.83 | 3.7E-02 | 1.69 | 0.53 | rna25181 |
| clu_3229_NA | rs43323 | -9.32 | 2.6E-05 | -0.96 | 0.81 | rna67556 |
| clu_3358_NA | rs18182 | -4.83 | 3.7E-02 | -0.91 | 0.53 | rna37159 |
| clu_3825_NA | rs17432 | -6.31 | 3.1E-03 | -3.87 | 0.65 | rna36077 |
| clu_4479_NA | rs7474 | 4.72 | 4.3E-02 | 0.99 | 0.51 | NA |
| clu_4676_NA | rs1560 | 5.78 | 7.6E-03 | 1.16 | 0.61 | NA |
| clu_470_NA | rs10737 | -4.89 | 3.3E-02 | -1.14 | 0.53 | rna22238 |
| clu_477_NA | rs10747 | 5.09 | 2.4E-02 | 1.00 | 0.55 | rna22297 |
| clu_5112_NA | rs23023 | 4.73 | 4.2E-02 | 1.08 | 0.52 | rna47697 |
| clu_5183_NA | rs23236 | 5.41 | 1.5E-02 | 0.97 | 0.58 | NA |
| clu_5274_NA | rs41125 | -11.43 | 1.8E-06 | -2.18 | 0.86 | NA |
| clu_5650_NA | rs3239 | -7.37 | 5.7E-04 | -2.12 | 0.72 | rna6239 |
| clu_5844_NA | rs3798 | 4.97 | 2.9E-02 | 1.12 | 0.54 | rna8159 |
| clu_5952_NA | rs9851 | 10.45 | 7.8E-06 | 1.52 | 0.84 | rna20451 |

| | | | | | | |
|---|---|---|---|---|---|---|
| clu_6005_NA | rs10060 | -12.85 | 3.0E-07 | -1.12 | 0.89 | rna20925 |
| clu_6148_NA | rs14722 | -9.65 | 1.7E-05 | -1.31 | 0.82 | NA |
| clu_6479_NA | rs400 | 4.64 | 5.0E-02 | 0.99 | 0.51 | rna628 |
| clu_6480_NA | rs401 | -4.80 | 3.8E-02 | -1.24 | 0.52 | rna628 |
| clu_6481_NA | rs401 | 5.57 | 1.1E-02 | 0.99 | 0.60 | rna630 |
| clu_6670_NA | rs37865 | 7.38 | 5.6E-04 | 1.16 | 0.72 | NA |
| clu_6976_NA | rs28636 | 9.20 | 3.2E-05 | 1.20 | 0.80 | rna56373 |
| clu_6978_NA | rs28627 | -13.89 | 7.4E-08 | -3.05 | 0.90 | rna56373 |
| clu_6980_NA | rs28608 | -11.26 | 2.3E-06 | -2.28 | 0.86 | rna56373 |
| clu_759_NA | rs33756 | 8.24 | 1.3E-04 | 2.89 | 0.76 | NA |
| clu_760_NA | rs33702 | 7.70 | 3.3E-04 | 1.74 | 0.74 | NA |

**Table S7 - *Cis*-eQTL associated with genes showing ecotype-associated expression patterns.** In some cases, several linked SNPs are associated with the same gene.

| Transcript | SNP | F-statistic | p-value | FDR | beta | $R^2$ | Gene symbol |
|---|---|---|---|---|---|---|---|
| rna14810 | rs7171 | 4.92 | 7.27E-05 | 0.04 | 0.62 | 0.54 | *TOMM5* |
| rna14810 | rs7175 | 4.75 | 1.08E-04 | 0.05 | 0.65 | 0.52 | *TOMM5* |
| rna1747 | rs813 | 5.13 | 4.36E-05 | 0.03 | 0.66 | 0.56 | *FAM83D* |
| rna1747 | rs815 | 4.94 | 6.86E-05 | 0.04 | 0.67 | 0.54 | *FAM83D* |
| rna19715 | rs9356 | -4.76 | 1.05E-04 | 0.05 | -0.94 | 0.52 | *TGM2* |
| rna19715 | rs9358 | -5.11 | 4.65E-05 | 0.03 | -0.76 | 0.55 | *TGM2* |
| rna19715 | rs9353 | -4.76 | 1.05E-04 | 0.05 | -0.94 | 0.52 | *TGM2* |
| rna19715 | rs9345 | 4.93 | 7.04E-05 | 0.04 | 0.79 | 0.54 | *TGM2* |
| rna19737 | rs9366 | 5.30 | 2.95E-05 | 0.03 | 0.54 | 0.57 | *SLC48A1* |
| rna2035 | rs901 | -5.37 | 2.52E-05 | 0.03 | -0.91 | 0.58 | *HESX1* |
| rna2035 | rs900 | -6.97 | 6.92E-07 | 0.01 | -0.95 | 0.70 | *HESX1* |
| rna2035 | rs898 | -5.37 | 2.52E-05 | 0.03 | -0.91 | 0.58 | *HESX1* |
| rna22829 | rs10950 | 7.94 | 9.30E-08 | 0.00 | 2.10 | 0.75 | *XMRK* |
| rna26480 | rs12534 | 4.95 | 6.69E-05 | 0.04 | 0.39 | 0.54 | *ANKRD24* |
| rna26480 | rs12533 | 4.95 | 6.69E-05 | 0.04 | 0.39 | 0.54 | *ANKRD24* |
| rna30313 | rs13850 | 5.12 | 4.53E-05 | 0.03 | 0.50 | 0.56 | *SDHAF4* |
| rna33863 | rs16144 | 4.88 | 7.90E-05 | 0.04 | 0.63 | 0.53 | *DPYSL3* |
| rna36030 | rs17316 | -6.43 | 2.24E-06 | 0.01 | -1.62 | 0.66 | *TBC1D10B* |
| rna36748 | rs17928 | -4.74 | 1.12E-04 | 0.05 | -0.64 | 0.52 | *CHP2* |
| rna37079 | rs18133 | 6.13 | 4.41E-06 | 0.01 | 0.51 | 0.64 | *SORL1* |
| rna37079 | rs18132 | 4.96 | 6.63E-05 | 0.04 | 0.40 | 0.54 | *SORL1* |
| rna38323 | rs18499 | 5.13 | 4.41E-05 | 0.03 | 0.98 | 0.56 | *PGK* |
| rna39492 | rs18912 | -5.20 | 3.71E-05 | 0.03 | -0.58 | 0.56 | *SOX19A* |
| rna4271 | rs2271 | 4.84 | 8.83E-05 | 0.04 | 0.90 | 0.53 | *NEDD8* |
| rna48702 | rs23575 | 4.89 | 7.77E-05 | 0.04 | 1.19 | 0.53 | *CCNG2* |
| rna48702 | rs23576 | 4.89 | 7.77E-05 | 0.04 | 1.19 | 0.53 | *CCNG2* |
| rna49634 | rs23831 | -6.16 | 4.10E-06 | 0.01 | -0.89 | 0.64 | *MRPS27* |
| rna50036 | rs24129 | -4.88 | 8.04E-05 | 0.04 | -0.67 | 0.53 | *U2AF2* |
| rna55686 | rs27912 | 5.06 | 5.19E-05 | 0.04 | 0.48 | 0.55 | *COMTD1* |

| | | | | | | | |
|---|---|---|---|---|---|---|---|
| rna57109 | rs29552 | -4.98 | 6.24E-05 | 0.04 | -0.35 | 0.54 | *TMEM106B* |
| rna62140 | rs35720 | 5.04 | 5.45E-05 | 0.04 | 0.48 | 0.55 | *BBIP1* |
| rna6601 | rs3380 | 6.98 | 6.77E-07 | 0.01 | 3.04 | 0.70 | *NEFM* |

**Table S8 - Transcripts with shared signatures of selection.**

| Transcript | Chromosome | Start | End | Gene symbol |
|---|---|---|---|---|
| rna11306 | nc_036845.1 | 5840150 | 5888150 | *LIMD1* |
| rna11655 | nc_036845.1 | 13274814 | 13292711 | *NR2F5* |
| rna11656 | nc_036845.1 | 13274814 | 13292711 | *NR2F5* |
| rna20015 | nc_036852.1 | 2440123 | 2450557 | *MSL3* |
| rna20016 | nc_036852.1 | 2440123 | 2449931 | *MSL3* |
| rna30685 | nc_036858.1 | 46729434 | 46752445 | *GALM* |
| rna38493 | nc_036863.1 | 25631283 | 25689381 | *NRIP1* |
| rna48244 | nc_036871.1 | 35450636 | 35453019 | *TIMM10* |
| rna54490 | nw_019942655.1* | 295040 | 302399 | *CD68* |
| rna43686 | nc_036867.1 | 38315700 | 38326913 | Uncharacterized LOC111953385 |
| rna49774 | nc_036873.1 | 650129 | 720173 | *ARHGAP10* |
| rna51987 | nc_036875.1 | 38852864 | 38915004 | *AHR* |
| rna17088 | nc_036849.1 | 20708415 | 20715252 | *KLHL28* |
| rna21979 | nc_036854.1 | 2386742 | 2392935 | *FAM49A 2C* |
| rna30924 | nc_036858.1 | 54659065 | 54673018 | *SUPV3L1* |

Note: *unplaced scaffold.
